# Optical measurement of glutamate release robustly reports short-term plasticity at a fast central synapse

**DOI:** 10.1101/2022.11.27.518035

**Authors:** Paul Jakob Habakuk Hain, Tobias Moser

## Abstract

Recently developed fluorescent neurotransmitter indicators have enabled direct measurements of neurotransmitter in the synaptic cleft. Precise optical measurements of neurotransmitter release may be used to make inferences about presynaptic function independent of electrophysiological measurement techniques. Here, we express iGluSnFR, a genetically encoded glutamate reporter in mouse spiral ganglion neurons to compare electrophysiological and optical readouts of presynaptic function and short-term synaptic plasticity at the endbulb of Held synapse. We conclude that iGluSnFR robustly and approximately linearly reports glutamate release from the endbulb of Held during synaptic transmission and allows assessment of short-term plasticity during high-frequency train stimuli. Furthermore, we show that iGluSnFR expression slightly alters the time course of spontaneous postsynaptic currents, but is unlikely to impact measurements of evoked synchronous release of many synaptic vesicles.

## Introduction

Recently, the advent of genetically encoded optical sensors has significantly enhanced the opportunities of experimental neurophysiology. Despite their wide use in various synapses (***Borghuis et al., 2013***; ***Taschenberger et al., 2016***; ***Sakamoto et al., 2018***; ***Pichler and Lagnado, 2019***; ***Özçete and Moser, 2021***; ***Vevea et al., 2021***; ***Mendonça et al., 2022***), relatively little attention has been given to potential interference with physiological neurotransmitter signaling and the established electrophysiological measurements.

Currently available genetically encoded glutamate indicators (GEGIs) are based either on AMPA receptors (AMPAR), like EOS (***Namiki et al., 2007***), or on bacterial glutamate binding proteins, such as GluSnFR, SuperGluSnFR (***Hires et al., 2008***), FLIPE (***Okumoto et al., 2005***), iGluSnFR (***Marvin et al., 2013***) and variants (***Marvin et al., 2018***; ***Helassa et al., 2018***). These optical methods allow highly resolved measurement of synaptic glutamate and its spatial profile in near-physiological conditions, but are limited in the temporal domain by their slow decay kinetics, and constraints set by photobleaching and lateral tissue movement (***Dürst et al., 2019***).

Yet they have helped overcome several problem of previously used techniques such as microdialysis (***Benveniste et al., 1984***) or enzymatically coupled amperometry (***Hu et al., 1994***), which have usually sampled over time frames of at least a few hundred milliseconds, too long to measure millisecond changes during synaptic activity.

On short time scales, presynaptic function is typically evaluated using electrophysiological measurements: either by presynaptic capacitance recordings, which measure the change in presynaptic membrane surface area during synaptic vesicle (SV) exocytosis (and endocytosis), or postsynaptic recordings, in which the presynaptic activity is filtered through the response characteristics of the postsynaptic receptors.

Both iGluSnFR and electrophysiological measurements give quantitative insights into the dynamics of glutamate release. Recently, ***Armbruster et al. (2020)*** indicated that iGluSnFR expression in synaptic or perisynaptic membranes can alter physiological glutamate signaling. They used Monte-Carlo simulations and recordings of astrocyte glutamate transporter currents in order to study the influence of iGluSnFR on glutamate dynamics by glutamate buffering. Their results suggest that iGluSnFR reduces the amount of free glutamate in and around the synaptic cleft, dependent on iGluSnFR expression levels and distance from the release site. Since the utility of GEGIs for the study of synaptic physiology relies on GEGIs or the expression system not interfering with physiological glutamate signaling, further experimental exploration of this proposed mechanism is warranted.

In the present study, we express iGluSnFR in the presynaptic membrane of auditory nerve fibers (ANFs) in the anteroventral cochlear nucleus (AVCN) to further investigate the potential influence on physiological glutamate signaling. Spherical and globular bushy cells (BCs) in the AVCN receive few large axo-somatic inputs from spiral ganglion neurons (SGNs) (***Liberman, 1993***; ***Tolbert and Morest, 1982***; ***Spirou et al., 2005***; ***Brawer and Morest, 1975***; ***Ryugo and Sento, 1991***; ***Cao and Oertel, 2010***). These large calyceal terminals are called endbulbs of Held (for the ANF – spherical BC synapse, ***Held (1893)***; ***Ryugo and Fekete (1982)***) or modified endbulbs of Held (ANF – globular BC, ***Rouiller et al. (1986)***), and are responsible for eliciting large, precisely timed, mainly AMPAR-mediated (***Antunes et al., 2020***; ***Cao and Oertel, 2010***; ***Sugden et al., 2002***; ***Schmid et al., 2001***; ***Gardner et al., 2001***; ***Wang et al., 1998***), EPSCs in the postsynaptic BC soma. Endbulbs of Held are a well-studied model synapses, where a wealth of details about the presynaptic function is known, mostly through the availability of highly resolved electrophysiological data (***Lin et al., 2011***; ***Cao and Oertel, 2010***; ***Yang and Xu-Friedman, 2012***, ***2009***, ***2008***; ***Wang and Manis, 2008***; ***Butola et al., 2017***).

Our combined iGluSnFR imaging and electrophysiological recordings suggest that iGluSnFR expression may slightly influence glutamate signalling. More specifically, we find changes in small excitatory postsynaptic currents elicited by spontaneous release of individual synaptic vesicles, but not in excitatory postsynaptic currents evoked by afferent fiber stimulation. Furthermore, we show that iGluSnFR signal behaves approximately as a linear transformation of postsynaptic current under physiological conditions, confirming theoretical predictions (***Armbruster et al., 2020***). We conclude that iGluSnFR is well-suited to measure some aspects of synaptic plasticity, but that its utility is limited by altering quantal release which it largely failed to resolve at the endbulb synapse.

## Results

### iGluSnFR expressed in SGNs reports glutamate release at the endbulb of Held

To measure glutamate release at the endbulb terminals of ANFs, we introduced iGluSnFR into SGNs. To our knowledge, no previous studies have used genetically encoded activity indicators in the cochlear nucleus. Here, a viral vector, adeno-associated virus (AAV) 2/9 carrying iGluSnFR under the control of the human synapsin promoter, was injected into the cochlea, targeting SGN cell bodies located in the modiolus. Injections through the round window are established in intracochlear pharmaco and gene therapy (e.g. ***Akil et al. (2012)***; ***Chen et al. (2003)***; ***Jung et al. (2015)***; ***Askew et al. (2015)***) and have been used to deliver small compounds like calcium dyes (***Chanda et al., 2011***; ***Zhuang et al., 2020***), optogenetic tools, including iGluSnFR (e.g. ***Keppeler et al. (2018)***; ***Özçete and Moser (2021)***), to SGN cell bodies and axons projecting onto inner hair cells.

To confirm successful viral gene transfer into SGNs, we first used immunohistochemistry to show the presence of iGluSnFR in the cochlea (fig. supp. 1 – 1, 2) and the cochlear nucleus (fig. 1, **A**). Next, we combined optical recordings with synchronous electrophysiological measurements, by patch-clamping BCs of acute brainstem slices and holding them at -70 mV, while evoking excitatory postsynaptic currents (eEPSCs) via monosynaptic input by monopolar stimulation through a saline filled pipette (fig. supp. 1 – 3). Afferent fiber stimulation is well-established in the cochlear nucleus(***Cao and Oertel, 2010***; ***Xu-Friedman and Regehr, 2005***) and allows one to elicit monosynaptic eEPSCs in BCs.

**Figure 1.**
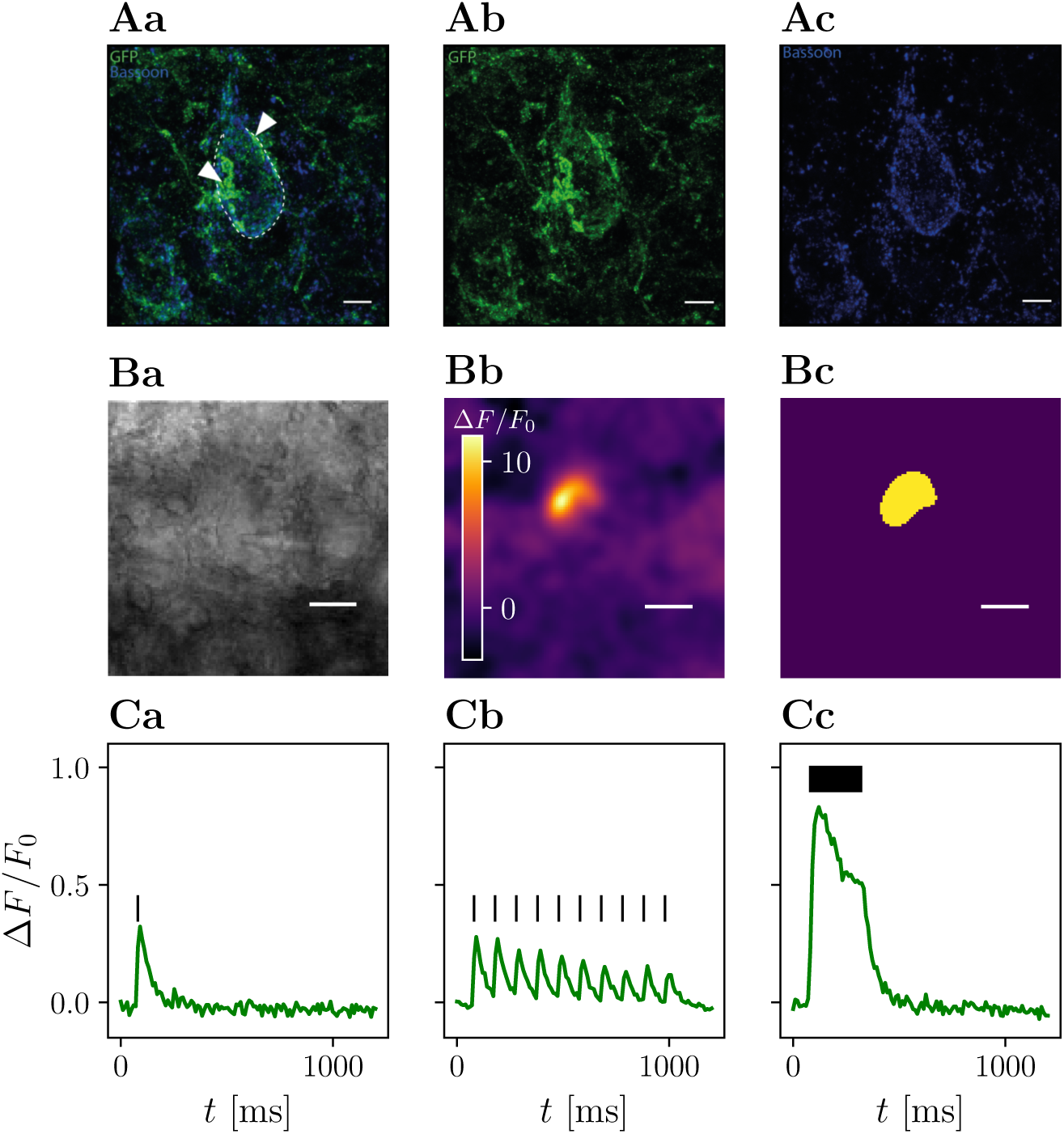
High-frequency stimulation of iGluSnFR-expressing presynaptic terminals allows discrimination of contiguous loci of glutamate release. Row A shows maximum-intensity projections immunohistochemical stainings of a BC in the AVCN. iGluSnFR expression was confirmed by staining for GFP (green), while Bassoon (blue) was used as a presynaptic marker. Panel Aa shows the overlay of both GFP and Bassoon stainings, while their isolated channels are presented in Ab and Ac, respectively. The central BC (dashed line) can be clearly differentiated by its one prominent protrusion towards the top of the cell, which differentiates BC from multipolar stellate cell. By their GFP staining, two putative endbulbs (arrow heads) can be identified towards the left and the right of the BC. Row B shows a representative example of the response of a single cell. In panel Ba a brightfield image of a patched bushy cell is shown. The Δ*F* image of three averaged responses to a repetitive stimulation of 25 stimuli at 100 Hz of the same cell in panel Bb shows a fluorescent response in a destinct cup-shaped area. A semi-automatic histogram-based segmentation algorithm selects a satisfactory ROI, giving an outer bound for the extent of the presynaptic terminal, shown in Bc. In row C, the quantified fluorescent responses of the same cell as in B to different stimulation paradigms is plotted over time. Plotting Δ*F*∕*F*_0_ over time in Ca, (average of 3 recordings) the response to a single stimulation (vertical bar) can be clearly identified. Stimulating at a 10 Hz frequency (10 stimuli) yields a fluorescent response average of 10 recordings), which clearly shows differentiable peaks (shown in Cb), which slightly overlap. The fluorescent response (average of 3 recordings) to repetitive stimulation (25 stimuli at 100 Hz) rises quickly in the first ∼ 100 ms, then decays to a plateau and after the last stimulation decays approximately exponentially back to the baseline, shown in Cc. Scale bars represent 5 µm. Figure 1—figure supplement 1. Immunohistochemistry of the cochlea I Figure 1—figure supplement 2. Immunohistochemistry of the cochlea II Figure 1—figure supplement 3. Bright-field example of a typical recording situation Figure 1—figure supplement 4. Comparison of ROI identification at different stimulation paradigms

We then compared different stimulation paradigms to find the maximum spatial extent of the presynaptic increase of iGluSnFR fluorescence (fig. supp. 1 – 4). The response to high-frequency stimulation best corresponded to the presumed terminal morphology and allowed efficient automatic image segmentation. We used the ROI extracted from the liberally filtered (Gaussian filter, σ = 3) Δ*F* image generated from recordings with 25 stimuli delivered at 100 Hz to establish an outer bound of the stimulated presynaptic terminal (fig. 1, **B**). Next, we plotted the iGluSnFR Δ*F*/*F*_0_ (measured as the change in fluorescence divided by the baseline fluorescence) response over time in the respective ROI (fig. 1, **C**) and found a large fluorescence increase in response to repetitive stimulation at 100 Hz. Analyzing the recordings for single stimuli and 10 Hz train stimulation, using the same ROI, showed a smaller signal (with a clear decay between stimulations for the 10 Hz train stimulation). Thus, we conclude that iGluSnFR expressed in SGNs via intracochlear round window injections of a viral vector is a suitable strategy to optically measure glutamate release dynamics in the cochlear nucleus.

### iGluSnFR expression prolongs the time course of mEPSCs

To address putative effects of iGluSnFR expression on synaptic transmission, we first analyzed spontaneous synaptic events at the endbulb of Held. In the endbulb of Held synapse, spontaneous excitatory postsynaptic currents (sEPSCs) in acute brainstem slices are not affected by tetrodotoxin (***Oleskevich and Walmsley, 2002***). We equate them to AMPAR-mediated miniature excitatory postsynaptic currents (mEPSCs) and tested whether they are affected by potential perturbations caused by the expression of iGluSnFR in the presynaptic SGN.

Representative mEPSC recordings from BCs, which were targeted by at least one iGluSnFR expressing endbulb and control BCs are displayed in fig. 2, **A**. iGluSnFR expression was confirmed by a subsequent fluorescent response to monopolar stimulation, while control mEPSC recordings were made from uninjected littermates of the AAV-injected mice. Fig. 2, **B** shows average mEPSC waves of *N* = 13 animals, *n* = 17 cells in the injected group and *N* = 11 control animals (*n* = 23 cells) and the kinetic parameters of the mEPSCs are presented in fig. 2, **C** tab. 1. While the mEPSC frequency, amplitude, and charge did not differ significantly, mEPSCs of BCs which were targeted by iGluSnFR-expressing ANFs decayed slightly slower (τ_decay_ = 0.182 ± 0.016 ms) than mEPSCs of control cells (τ_decay_ = 0.145 ± 0.007 ms, *p* = 0.003) and were slightly wider (full width at half maximum (FWHM) = 0.251 ± 0.011 ms) than control cells (FWHM = 0.214 ± 0.007 ms, *p* = 0.004).

**Figure 2.**
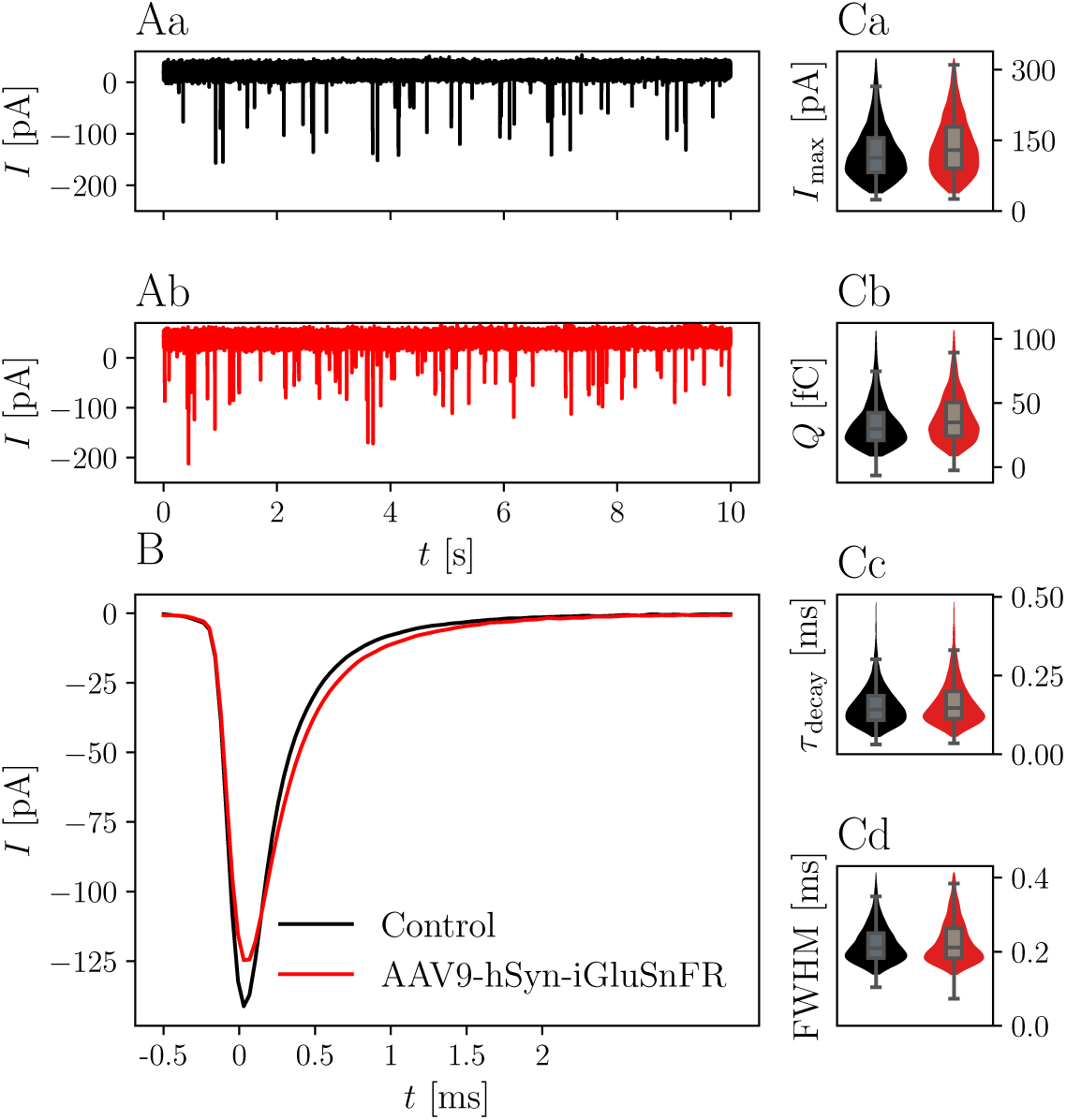
iGluSnFR expression in the presynaptic membrane prolongs the mEPSC decay time without hanging the overall glutamate charge. Panel Aa shows a representative 10 s trace of a control BC held at -70 mV. Panel Ab shows a corresponding trace of a BC which was targeted by at least one iGluSnFR-expressing SGN. Panel B shows overlayed the average peak-aligned mEPSC for *N* = 13 animals and *n* = 17 cells in the iGluSnFR condition and *N* = 11 animals, *n* = 23 cells in the control condition. Panels C show violin plots with overlayed boxplots of the distribution of parameters of mEPSCs over all cells. The parameters are mEPSC amplitude (*I*_max_), charge (*Q*), time constant of a single exponential fitted through the decay τ_decay_ and full-width at half maximum (FWHM). For illustration purposes, violin plots were constrained to values within the 0.1 – 0.99 interquantile range. Boxes indicate the median, the 0.25 and 0.75 quantiles with whiskers spanning 1.5 times in interquartile range (outliers not shown).

**Table 1.** Measured and derived parameters of mEPSCs

|  | iGluSnFR | Control | *p* |
| --- | --- | --- | --- |
| $\alpha^* = 0.010$ | | | |
| *N* | 13 | 11 |  |
| *n* | 17 | 23 |  |
| Events | 7197 | 8707 |  |
| *f* [Hz] | 7.44 ± 2.00 | 6.33 ± 1.76 | 0.292<sup>†</sup> |
| *I*<sub>max</sub> [pA] | 126.7 ± 9.2 | 129.4 ± 7.9 | 0.732<sup>‡</sup> |
| *Q* [fC] | 39.1 ± 0.2 | 34.6 ± 0.2 | 0.218<sup>‡</sup> |
| $\tau_{\text{decay}}$ [ms] | 0.182 ± 0.016 | 0.145 ± 0.007 | 0.003<sup>‡</sup> |
| FWHM [ms] | 0.251 ± 0.011 | 0.214 ± 0.007 | 0.004<sup>‡</sup> |

Parameters of mEPSCs of bushy cells that were targeted by at least one iGluSnFR-expressing endbulb and bushy cells of control animals. For each cell, 60s of spontaneous activity was recorded. Data is presented as mean ± standard error of the mean. <sup>†</sup> Frequencies (*f*) were compared with a Wilcoxon rank sum test. <sup>‡</sup> *p*-values for the coefficients of a mixed linear effects model with a fixed treatment effect and a random cell-specific intercept for mEPSC amplitude (*I*<sub>max</sub>), charge (*Q*), decay time constant ( $\tau_{\text{decay}}$ ) and full-width at half maximum (FWHM). $\alpha^*$ is the Bonferroni-adjusted significance level.

Assuming AMPARs are not saturated by a single quantum of glutamate (***Ishikawa et al., 2002***; ***Yamashita et al., 2003***), an extracellular glutamate buffer, prolonging the time course in the synaptic cleft, while leaving the total amount of glutamate released unaffected, would alter the time profile of the postsynaptic currents, but not the integrated charge. According to this hypothesis, we expect the peak current to be reduced under the influence of iGluSnFR. This, however, was not observed to a statistically significant degree.

A caveat of our analysis is that mEPSCs recorded in a given BC may originate from synapses that vary in iGluSnFR expression. During some experiments on slices of AAV-injected animals, BCs with large eEPSCs failed to show any iGluSnFR fluorescence increase (data not shown). Even though these recordings were not included in the analysis of evoked release, their presence highlights the possibility that of the multiple endbulbs projecting onto the same BC, some did not express iGluSnFR. Since we ensured single-fiber stimulation and excluded recordings without iGluSnFR fluorescence increase, iGluSnFR-negative endbulbs did not contribute to the eEPSCs we measured. Yet, spontaneous release from iGluSnFR-negative endbulbs is expected, hence only a fraction of the measured quantal events originated from iGluSnFR-expressing endbulbs. Consequently, the differences may be a lower bound of the effect of iGluSnFR expression on single synapses.

### iGluSnFR expression does not alter evoked EPSCs

Given the effect of iGluSnFR expression on quantal currents, we further investigated possible effects on synaptic transmission by analyzing evoked release. In the endbulb of Held, monopolar stimulation is able to induce eEPSCs in the nA range, corresponding to the synchronized release of several tens to hundreds of SVs. Comparing the response of BCs postsynaptic to iGluSnFR-expressing and control endbulbs to a single electrical stimulus, we did not find significant differences in amplitude and kinetics (summarized in tab. 2).

**Table 2.** Measured and derived parameters of eEPSCs

|  | iGluSnFR | Control | *p* |
| --- | --- | --- | --- |
| $\alpha^* = 0.004$ | | | |
| 2mM [Ca<sup>2+</sup>]<sub>e</sub> |  |  |  |
| *N* | 7 | 7 |  |
| *n* | 8 | 13 |  |
| *t*<sub>syn delay</sub> [ms] | 0.935 ± 0.054 | 0.890 ± 0.029 | 0.284 |
| *I*<sub>max</sub> [nA] | 1.74 ± 0.32 | 1.41 ± 0.19 | 0.241 |
| *Q* [pC] | 0.94 ± 0.18 | 0.57 ± 0.1 | 0.039 |
| *t*<sub>rise</sub> [ms] | 0.189 ± 0.004 | 0.168 ± 0.007 | 0.640 |
| $\tau_{\text{decay}}$ [ms] | 0.334 ± 0.031 | 0.305 ± 0.015 | 0.392 |
| FWHM [ms] | 0.429 ± 0.042 | 0.410 ± 0.026 | 0.544 |
| 4mM [Ca<sup>2+</sup>]<sub>e</sub> |  |  |  |
| *N* | 7 | 3 |  |
| *n* | 9 | 7 |  |
| *t*<sub>syn delay</sub> [ms] | 0.960 ± 0.047 | 1.064 ± 0.044 | 0.088 |
| *I*<sub>max</sub> [nA] | 4.85 ± 1.03 | 6.01 ± 1.75 | 0.589 |
| *Q* [pC] | 3.01 ± 0.73 | 5.75 ± 2.59 | 0.297 |
| *t*<sub>rise</sub> [ms] | 0.226 ± 0.035 | 0.210 ± 0.015 | 0.295 |
| $\tau_{\text{decay}}$ [ms] | 0.327 ± 0.037 | 0.382 ± 0.056 | 0.288 |
| FWHM [ms] | 0.523 ± 0.035 | 0.543 ± 0.079 | 0.846 |

Parameters of eEPSCs of iGluSnFR-expressing endbulb synapses and endbulb synapses of control animals at 2 and 4 mM [Ca<sup>2+</sup>]<sub>e</sub>. *p*-values for the coefficients of a mixed linear effects model with a fixed treatment effect and a random cell-specific intercept for synaptic delay (*t*<sub>syn delay</sub>), eEPSC amplitude (*I*<sub>max</sub>), charge (*Q*), rise time (*t*<sub>rise</sub>), decay time constant ( $\tau_{\text{decay}}$ ) and full-width at half maximum (FWHM). $\alpha^*$ is the Bonferroni-adjusted significance level.

At 2 mM [Ca^2+^]_*e*_ we compared eEPSCS in BCs (*n* = 8) of AAV-injected mice (*N* = 7) to those of uninjected littermates (*N* = 7, *n* = 13). While the amplitude of eEPSCs was increased, the difference was not statistically significant (*I*_max, iGluSnFR_ = 1.74 ± 0.32 nA, *I*_max, Control_ = 1.41± 0.19 nA, *p* = 0.241 at *α*^∗^ = 0.004).

We then measured eEPSCs at 4 mM [Ca^2+^]_*e*_ (AAV-injected group: *N* = 7, *n* = 9, control group: *N* = 3, *n* = 7). The EPSC size in the AAV-injected group increased from 1.74 ± 0.32 nA at 2 mM [Ca^2+^]_*e*_ to 4.85 ± 1.03 nA at 4 mM [Ca^2+^]_*e*_ (*p* < 0.001). Likewise, in the control group, eEPSC sizes increased from 1.41 ± 0.19 nA to 6.01 ± 1.75 nA (*p* < 0.001). Comparison at 4 mM [Ca^2+^]_*e*_, did not reveal statistically significant differences (*I*_max, iGluSnFR_ = 4.85 ± 1.03 nA, *I*_max, Control_ = 6.01 ± 1.75 nA, *p* = 0.589). Neither at 2 mM [Ca^2+^]_*e*_, nor at 4 mM [Ca^2+^]_*e*_, we saw a statistically significant change in charge, FWHM, rise time, decay time constant and synaptic delay (see tab. 2).

We then analyzed the eEPSCs to repetitive stimulation in the same iGluSnFR-expressing and control endbulbs. First, we considered the paired-pulse ratio (PPR), i.e. the ratio of two consecutive EPSCs with a defined inter-stimulus interval (ISI), presented in fig. 3. Comparing the PPRs with ISIs of 10 ms and 100 ms (see fig. 3 and tab. 3) in 2 mM [Ca^2+^]_*e*_ conditions revealed a statistically significant increase in the PPR_10 ms_ (PPR_10 ms, Control_ = 0.82 ± 0.05 and PPR_10 ms, iGluSnFR_ = 1.00 ± 0.05 *p* = 0.004), which was absent at a longer ISI.

**Figure 3.**
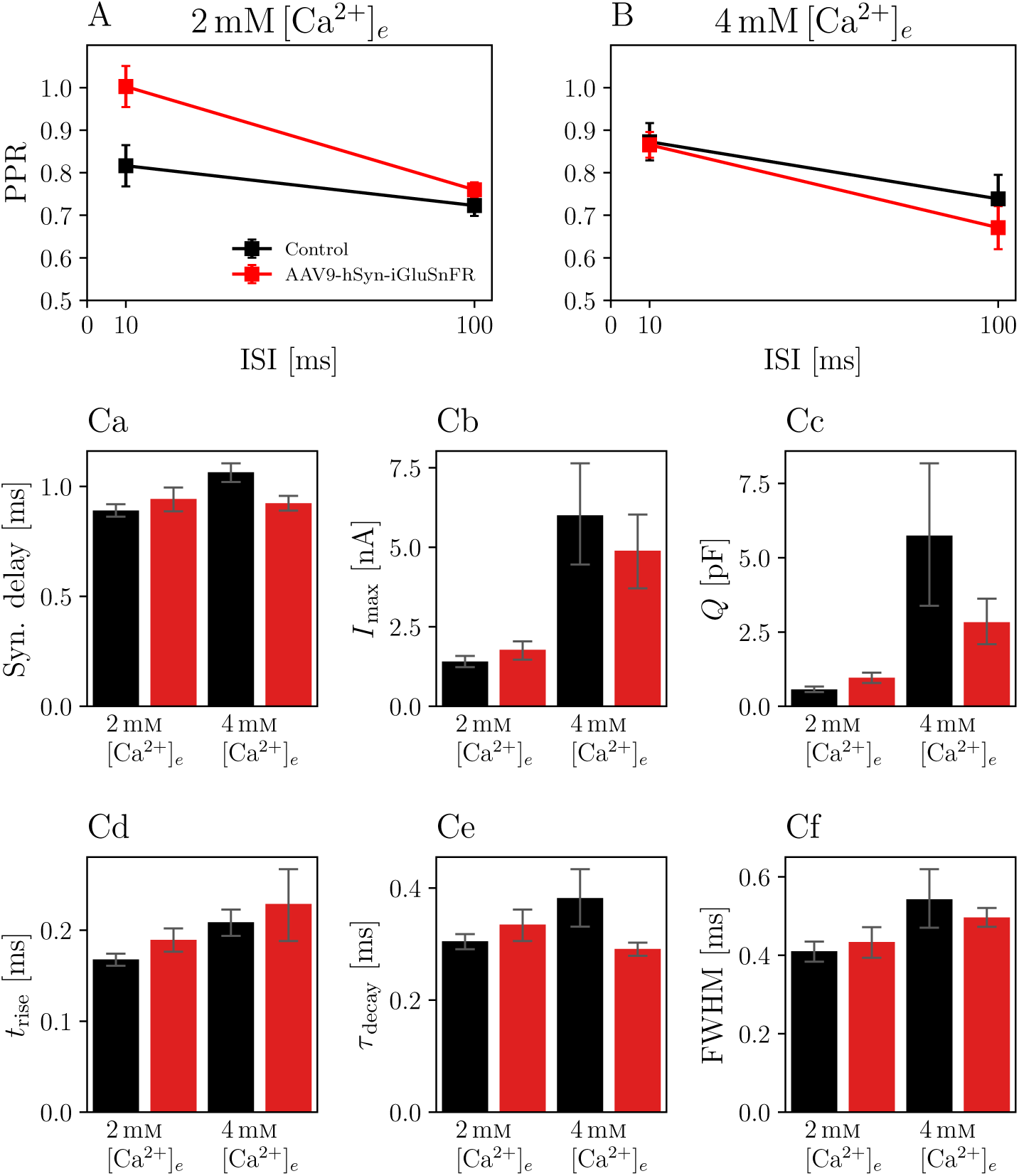
Paired pulse ratio (PPR) of iGluSnFR-expressing endbulb synapses is increased for 10 ms interstimulus interval (ISI) at 2 mM [Ca^2+^]_*e*_, but remains unchanged at higher calcium concentrations or longer ISIs. Plots show the PPR for ISI of 10 ms (PPR_10 ms_) and 100 ms (PPR_100 ms_) of *N* = 7 animals, *n* = 8 cells in the iGluSnFR group and *N* = 7 animals, *n* = 13 cells in the control group under 2 mM [Ca^2+^]_*e*_ conditions (in A), and *N* = 7 animals, *n* = 9 cells in the iGluSnFR group and *N* = 3 animals, *n* = 7 cells in the control group at 4 mM [Ca^2+^]_*e*_ (in B). Panels C show bar plots with of kinetic parameters of EPSCs, namely synaptic delay Ca, eEPSC amplitude (*I*_max_, Cb), charge (*Q*, Cc), rise time (*t*_rise_, Cd), time constant of a single exponential fit through the decay phase of the eEPSC (τ_decay_,Ce) and full-width at half maximum (FWHM, Cf). Bars represent means ± standard error of the mean of the median values per cell.

**Table 3.** Comparisons of PPRs

|  | iGluSnFR | Control | *p* |
| --- | --- | --- | --- |
| $\alpha^* = 0.013$ | | | |
| 2 mM $[\text{Ca}^{2+}]_e$ | | | |
| *N* | 7 | 7 |  |
| *n* | 8 | 13 |  |
| PPR<sub>10ms</sub> | 1.00 ± 0.05 | 0.82 ± 0.05 | 0.004 |
| PPR<sub>100ms</sub> | 0.76 ± 0.02 | 0.72 ± 0.02 | 0.114 |
| 4 mM $[\text{Ca}^{2+}]_e$ | | | |
| *N* | 7 | 3 |  |
| *n* | 9 | 7 |  |
| PPR<sub>10ms</sub> | 0.87 ± 0.03 | 0.87 ± 0.04 | 0.477 |
| PPR<sub>100ms</sub> | 0.67 ± 0.05 | 0.74 ± 0.06 | 0.059 |

Paired pulse ratios (PPR) for 10 ms and 100 ms inter-stimulus intervals for iGluSnFR-expressing endbulbs and control synapses at 2 and 4 mM $[\text{Ca}^{2+}]_e$ . *p*-values are given for the coefficients of a mixed linear effects model with a fixed treatment effect and a random cell-specific intercept. $\alpha^*$ is the Bonferroni-adjusted significance level.

In order to assess potential differences in presynaptic activity, we further analyzed data collected from stimulus trains. If the expression of a transmembrane protein in the presynaptic membrane had an effect on the release of SVs, standard estimates of vesicle pool parameters might reflect these perturbations. Furthermore, if iGluSnFR was saturated with glutamate within the first few stimuli and stayed saturated for the remaining train, the current elicited by one SV in the beginning of a train, might be different from the current elicited by one SV in the end of the train, leading to differences in standard cumulative plots used to analyze high-frequency train data (***Schneggenburger et al., 1999***).

In fig. 4, recordings of 25 stimuli at 100 Hz are shown. The averaged raw traces (plots in row A) and current amplitudes (row B) show the characteristic increase of release at elevated [Ca^2+^]_*e*_ concentrations and the increase in PPR_10 ms_ at 2 mM [Ca^2+^]_*e*_. There was no significant change in pool parameters or release probability derived from the cumulative plots, summarized in tab. 4).

**Figure 4.**
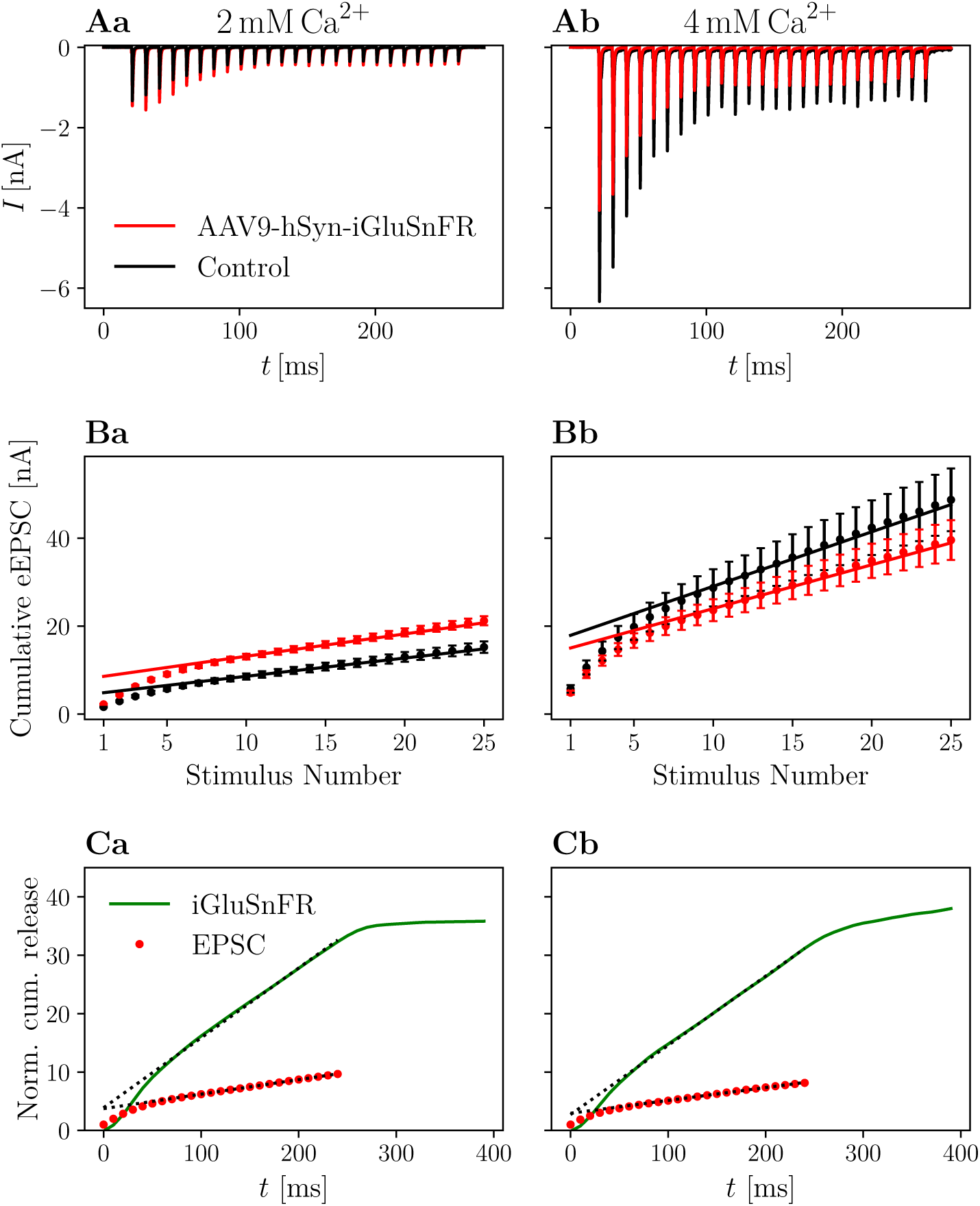
Cumulative release is not significantly altered in iGluSnFR-expressing synapses and cumulative release analysis of iGluSnFR data is comparable to standard electrophysiological approaches. Panel A shows the absolute amplitude of eEPSCs elicited by 25 stimuli at 100 Hz, while in B the cumulative EPSC amplitude is plotted over the stimulus number in order to obtain estimates for RRP size, release probability and replenishing rate (see ***Schneggenburger et al. (1999)***; ***Neher (2015)***). In C, cumulative release derived from iGluSnFR recordings and eEPSC recordings from injected animals are plotted over time. The dashed line shows the fit of the steady state response, continued to the ordinate. Error bars omitted for clarity. In a the data recorded from *N* = 7 animals, *n* = 8 cells in the iGluSnFR group and *N* = 7 animals, *n* = 13 cells in the control group (in panels A, B) under 2 mM [Ca^2+^]_*e*_ conditions is shown. In b, data recorded from *N* = 7 animals, *n* = 9 cells in the iGluSnFR group and *N* = 3 animals, *n* = 7 cells in the control group (in panels A, B) at 4 mM [Ca^2+^]_*e*_ is shown. Figure 4—figure supplement 1. Cumulative release analysis of 10 Hz data

**Table 4.** Pool parameters derived from cumulative eEPSCs

|  | iGluSnFR | Control | *p* |
| --- | --- | --- | --- |
| $\alpha^* = 0.008$ | | | |
| 2 mM $[\text{Ca}^{2+}]_e$ | | | |
| *N* | 7 | 7 |  |
| *n* | 8 | 13 |  |
| RRP [nA] | 8.455 ± 1.608 | 5.137 ± 0.882 | 0.060 |
| $r_{\text{refill}}$ [nA ms<sup>-1</sup>] | 0.053 ± 0.006 | 0.042 ± 0.003 | 0.248 |
| $p_r$ | 0.27 ± 0.02 | 0.34 ± 0.02 | 0.009 |
| 4 mM $[\text{Ca}^{2+}]_e$ | | | |
| *N* | 7 | 3 |  |
| *n* | 9 | 7 |  |
| RRP [nA] | 15.051 ± 2.210 | 21.372 ± 5.750 | 0.300 |
| $r_{\text{refill}}$ [nA ms<sup>-1</sup>] | 0.097 ± 0.021 | 0.138 ± 0.035 | 0.310 |
| $p_r$ | 0.32 ± 0.02 | 0.31 ± 0.02 | 0.904 |

Pool parameters for iGluSnFR-expressing endbulbs and control synapses at 2 and 4 mM $[\text{Ca}^{2+}]_e$ . *p*-values for the coefficients of a mixed linear effects model with a fixed treatment effect and a random cell-specific intercept were computed for the readily releasable pool (RRP) in nA, the refilling rate ( $r_{\text{RRP}}$ ) and release probability in resting conditions ( $p_r$ ) of iGluSnFR and control group were compared with a Wilcoxon rank sum test. $\alpha^*$ is the Bonferroni-adjusted significance level.

### Glutamate imaging yields quantitative measures of synaptic depression

To assess the feasibility of measuring evoked glutamate release with a GEGI and the validity of its use for estimating parameters of synaptic release, we systematically correlated iGluSnFR measurements with electrophysiological read-outs of glutamate release at 2 mM [Ca^2+^]_*e*_ (*N* = 7 animals, *n* = 8 cells) and at 4 mM [Ca^2+^]_*e*_ (*N* = 7, *n* = 9).

Since the signal-to-noise ratio (SNR) of iGluSnFR recordings was not sufficient to resolve the release of single SV in the preparation in our hands, we aimed for a robust protocol correlating electrophysiological and iGluSnFR recordings. This was achieved by measuring synaptic responses to single fiber stimulations both by fluorescence imaging at a frame rate of 100 Hz (fig. 5, **A, C**) and voltage-clamp recordings of eEPSCs (fig. 5, **B, D**). The peak Δ*F*∕*F*_0_ response amounted to 0.30 ± 0.02 (*N* = 7, *n* = 8). For comparison, the maximal Δ*F*∕*F*_0_ response of 4.5 has been reported to for iGluSnFR *in vitro*, and Δ*F*∕*F*_0_ values around 1 have been described in cells (***Marvin et al., 2013***), which were achieved using preparations with less background noise (i.e. dissociated cultured cells), stronger illumination and imaging techniques, such as laser scanning confocal microscopy, which enable less noisy image acquisition. A recent study using iGluSnFR for reporting glutamate release from hair cells of the organ of Corti (same viral delivery approach as in the present study) reported maximal Δ*F*∕*F*_0_ values of around 0.5 using spinning disk confocal imaging (***Özçete and Moser (2021)***, their figure EV1).

**Figure 5.**
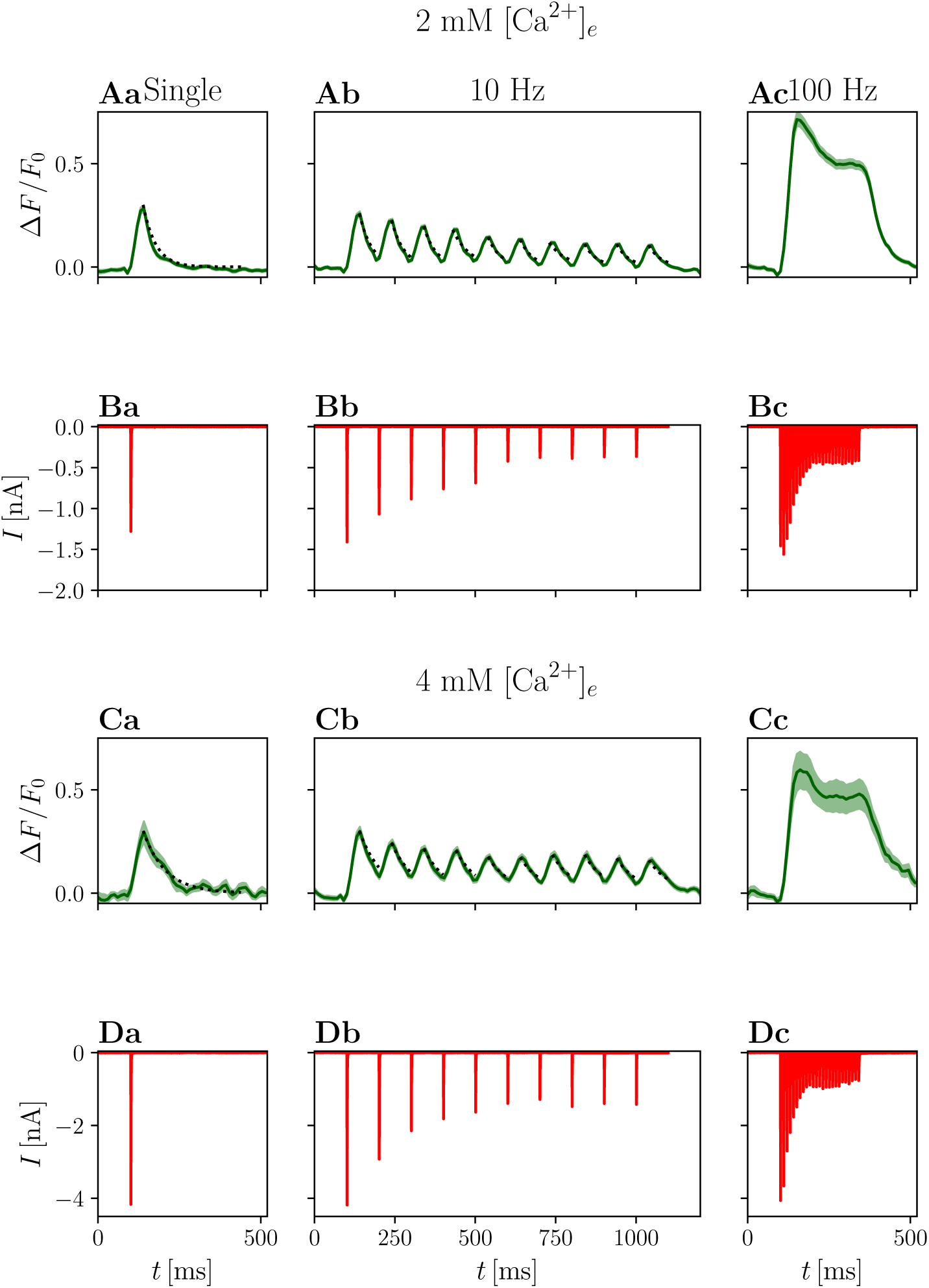
Electrophysiological and optical responses to different stimulation paradigms. Panel A shows grand averages of optical responses at 2 mM [Ca^2+^]_*e*_ from *N* = 7 animals, *n* = 8 cells and panel B shows the corresponding eEPSCs. Panels C and D show the respective data recorded at 4 mM [Ca^2+^]_*e*_ from *N* = 7 animals, *n* = 9 cells. In a, average responses to a single stimulus are presented, b shows the average response to 5 or 10 stimuli (pooled for presentation purposes) at 10 Hz and c to 25 stimuli at 100 Hz. In the plots of the optical responses to a single stimulation, an exponential fit through the decay is plotted as a dotted line. The same exponential function is plotted to the peaks of the corresponding 10 Hz data.

The decay of the signal was fit with a single exponential function: for a single stimulation in 2 mM [Ca^2+^]_*e*_, the average time constant τ_decay_ was 38.43 ± 0.01 ms (fig. 5, **A**). In elevated [Ca^2+^]_*e*_ conditions (4 mM, fig. 5, **C**), the mean peak Δ*F*∕*F*_0_ values was 0.32 ± 0.06, which is not significantly different from the values measured at 2 mM [Ca^2+^]_*e*_ conditions (*p* = 0.294). In contrast, τ_decay_ was nearly doubled (68.66 ± 0.01 ms, *p* < 0.001).

It can be assumed that significantly more glutamate is released in 4 mM [Ca^2+^]_*e*_ conditions, as EPSC size increased from 2.05 ± 0.06 nA to 4.83 ± 0.24 nA (*p* < 0.001) in the presence of kynurenate. This effect agrees well with the previously described phenomenon that during stimulation with multiple closely spaced pulses, adding more pulses prolongs the response, but does not increase the amplitude (***Marvin et al., 2013***). Similarly, decay times have been reported to increase with prolonged presynaptic depolarization in IHC (***Özçete and Moser, 2021***), suggesting that iGluSnFR response kinetics may depend on the amount of released glutamate or that it is saturated and returns from saturation more slowly if glutamate remains elevated for longer.

Further investigating the iGluSnFR response to evoked synaptic glutamate release, we stimulated the afferent ANF with trains of stimuli at 10 Hz, a frequency at which we expected the peaks of the iGluSnFR response to overlap minimally. Fig 5 shows that the τ_decay_ are approximately constant across stimulations, suggesting that there is no significant build-up of glutamate during this train of stimulation. Individual responses cannot be discriminated for stimulation trains of higher frequency, such as 25 stimuli at 100 Hz. Rather, the response consists of a large peak at the onset of the stimulation, which decays first to a plateau and, after the end of the stimulation, back to the baseline (fig. 5, **Ac, Cc**).

Next, we compared electrophysiological with optical readouts of presynaptic plasticity. Amplitudes of both eEPSC and iGluSnFR responses decline during a 10 Hz train (see fig. 5, **b**), as a consequence of presynaptic depression. As both AMPAR currents and iGluSnFR signals may relate non-linearly to the amount of released glutamate, we plot the normalized area under the curve of the Δ*F*∕*F*_0_ iGluSnFR signal for each pulse in a train of 5 or 10 stimuli at 10 Hz versus the corresponding normalized eEPSC amplitude. One limitation inherent to this analysis results from the partial overlap of subsequent iGluSnFR responses, an effect that is even more pronounced at 4 mM [Ca^2+^]_*e*_ with longer individual responses.

To ameliorate this problem, 10 Hz iGluSnFR responses were subjected to deconvolution analysis. The comparison of the normalized peaks of the deconvolved average response to a train of 5 or 10 stimuli at 10 Hz and the corresponding eEPSC magnitude is plotted in fig. 6, **C**. Comparing the different measures of glutamate release, fig. 6, **C** shows that the deconvolved iGluSnFR response and eEPSC are approximately linearly related, suggesting a similar relationship to the underlying presynaptic glutamate release. Supp. fig. 6 – 1 shows corresponding analysis of non-normalized data. Supp. fig. 6 – 2 shows raw deconvolved traces of iGluSnFR responses.

**Figure 6.**
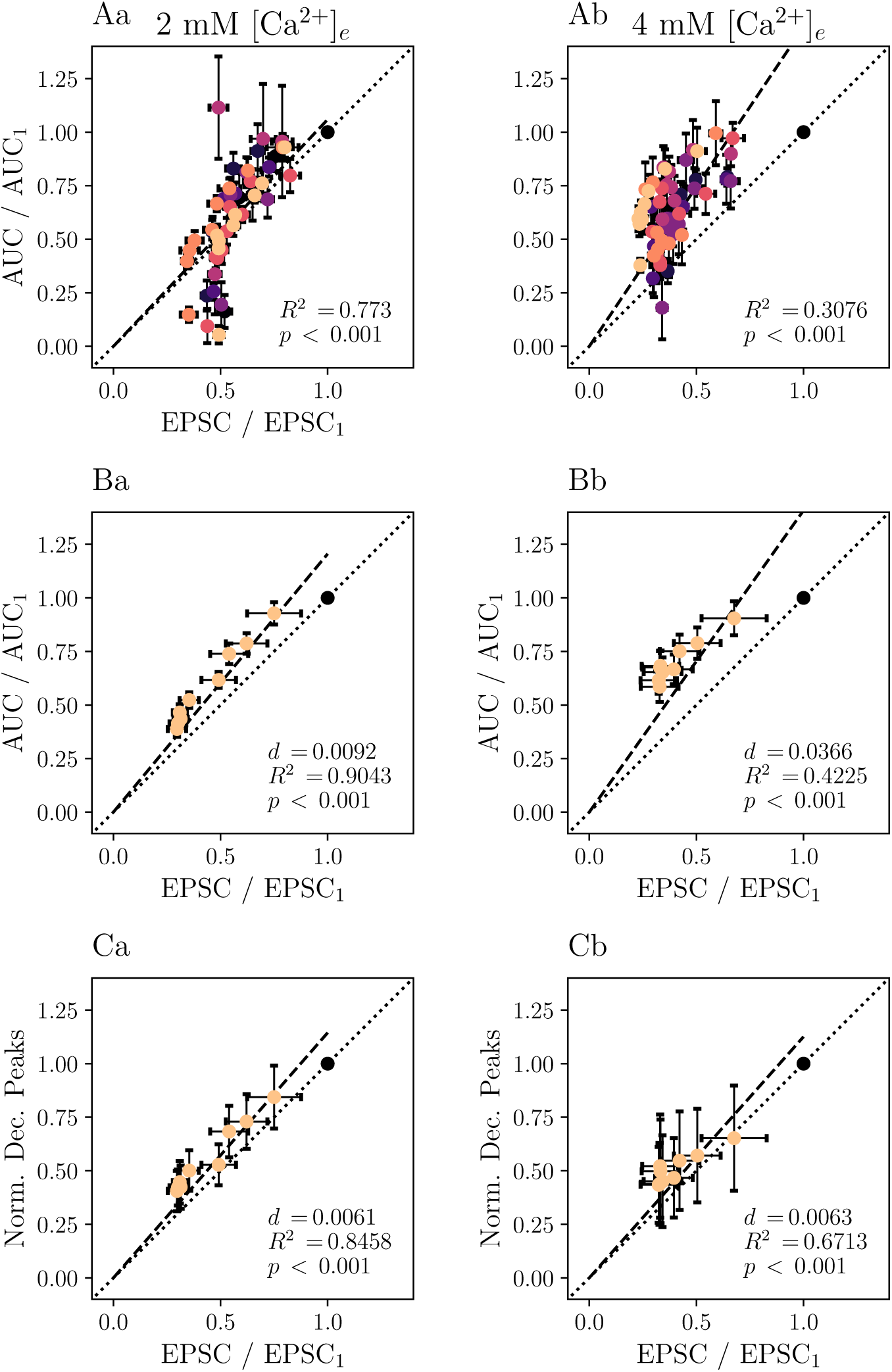
Deconvolution analysis improves the agreement of iGluSnR-derived and electrophysiologically derived measures of short-term plasticity. Plots based on recordings of of *N* = 7 animals, *n* = 8 cells performed in 2 mM [Ca^2+^]_*e*_ conditions are displayed in the a panels, those based on those of *N* = 7 animals, *n* = 9 cells at 4 mM [Ca^2+^]_*e*_ in panels b. If values for individual cells are shown, data points derived from the same cell are plotted in the same color. Values are derived from 10 to 20 recordings per cell. In panel A, the AUC normalized to the first of the iGluSnFR data is plotted against the respective normalized eEPSC amplitude for individual cells. Panel B shows grand averages of the data presented in A. In C, the peaks of the deconvolved iGluSnFR, normalized to the first stimulation, are plotted against normalized eEPSC. For the data in all panels, a linear regression model was applied (constrained to functions passing through the origin, dashed lines). Dotted lines correspond to the the linear function with the slope 1. For each regression model, coefficients of determination *R*^2^ and *p* values were calculated. For panels B and C, the average squared distances *d* of each point to the identity function is given. Figure 6—figure supplement 1. Non-normalized data Figure 6—figure supplement 2. Deconvolved traces of averaged recordings

The results presented in fig. 6 thus suggest that iGluSnFR shows little saturation, at least when compared to AMPAR-mediated eEPSCs in the presence of 1 mM kynurenate. A previous study showed that in IHCs, that the iGluSnFR Δ*F*∕*F*_0_ and its integral correlate well with changes in presynaptic membrane capacitance, another measure of presynaptic SV exocytosis (***Özçete and Moser, 2021***). Further exploring the viability of analyzing transmitter release by iGluSnFR imaging, we analyzed cumulative release rates as measured by iGluSnFR and compared them to values derived from the analysis of EPSCs. First, we analyzed trains at 10 Hz (fig. supp. 4 – 1). The average steady state iGluSnFR response (stimuli 5 – 10) was depressed to 0.33 times the amplitude of the first stimulus in 2 mM [Ca^2+^]_*e*_ and to 0.34 times the amplitude at 4 mM [Ca^2+^]_*e*_. We note that this moderate depression requires a correction for the analysis of cumulative release (***Thanawala and Regehr (2013)***, but also see ***Neher (2015)***) to accurately assess vesicle pool parameters, while accounting for incomplete RRP depletion. This correction would need to take into account the different release probabilities for the first event, *p*_*r*,1_, and the events at a steady state *p*_*r*,*ss*_ and would have to be applied to both imaging and electrophysiological data.

Since in the current approach, release probabilities, specifically later in the train, cannot be easily inferred, we chose to not apply the correction and instead report *y*_intercept_/*y*_1_, where *y*_1_ is the first response, *y*_intercept_ is the intercept with the ordinate of the affine function fit through the later responses and *p*_*r*_ = (*y*_intercept_/*y*_1_) ⋅ (*p*_*r*,1_/*p*_*r*,*ss*_). This approach still yields useful information on the comparability of the two approaches, as the error introduced by not correcting for incomplete depletion of the RRP is identical for iGluSnFR data and EPSC recordings. For stimulations at 10 Hz, *y*_intercept_/*y*_1_ was 0.52 at 2 mM [Ca^2+^]_*e*_ and 0.65 at 4 mM [Ca^2+^]_*e*_ for iGluSnFR recordings, while electrophysiological recordings yielded 0.44 at 2 mM [Ca^2+^]_*e*_ and 0.63 at 4 mM [Ca^2+^]_*e*_.

Even though iGluSnFR recordings of responses to trains of higher frequencies do not give access to individual peaks, deconvolution analysis has been used before to determine cumulative release rates (***Sakamoto et al., 2018***). In fig. 4, **C** the cumulative release rates, as measured by deconvolution analysis of responses to 100 Hz trains is juxtaposed to SMN plots. Calculating *y*_intercept_/*y*_1_ on the basis of deconvolved iGluSnFR recordings yielded 0.25 at 2 mM [Ca^2+^]_*e*_ and 0.37 at 4 mM [Ca^2+^]_*e*_. Performing an SMN analysis with recorded eEPSC gave 0.27 at 2 mM [Ca^2+^]_*e*_ and 0.34 at 4 mM [Ca^2+^]_*e*_. Parameters estimated from the imaging data compare favorably to data estimated from electrophysiological recordings, especially higher-frequency stimulation at 100 Hz. Nonetheless, it is important to note the limited physiological relevance of these estimates due to the comparatively small amount of stimulations and the associated problems (***Neher, 2015***). Specifically, parameters derived from recordings at 10 Hz should be interpreted with caution, as pool depletion is likely incomplete and the steady-state might not have been reached by the end of the train. These caveats apply to measured currents in the same way as to the iGluSnFR data, such that the results suggest that estimation of pool parameters from imaging data is viable, although subject to the same constraints as the estimation from EPSC data. Additionally, the size of the RRP cannot be estimated in terms of individual vesicles as we did not resolve the iGluSnFR signal corresponding to the release of individual vesicles.

## Discussion

Fluorescent glutamate indicators have significantly increased our ability to monitor presynaptic activity of glutamatergic synapses. In this study, we sought to address two important potential caveats of working with iGluSnFR. First, pertaining to the question if the presence of iGluSnFR alters physiological glutamate release, we show an effect on some, but not all postsynaptic currents. Small currents, elicited by single vesicles decay slower and are significantly broader, while we did not find a corresponding change in eEPSCs. Additionally, we find evidence, suggesting an impact on successive stimuli, at least at short intervals between stimuli. Secondly, we find an almost linear relationship of iGluSnFR signal to electrophysiological estimates of postsynaptic activity, at least when using deconvolution analysis.

### Monitoring glutamate release with optical sensors

On a single synapse level, direct measurement of neurotransmitter release by optical means allows for the dissection of pre and postsynaptic mechanisms of information processing, such as short term or homeostatic plasticity. Commonly available electrophysiological techniques for studying presynaptic activity yield either highly resolved recordings of single neurons or information about the activity of neuronal populations. Highly-resolved single cell techniques, such as intracellular recordings are usually limited in the number of cells that can be observed in parallel, while population-level recordings, such as by microelectrode arrays have shortcomings in identifying recorded neurons and depth of analysis.

Compared to more broadly employed imaging methods such as presynaptic Ca^2+^ imaging as a proxy for synaptic activity, direct measurement of presynaptic neurotransmitter release offers distinct advantages. First, presynaptic Ca^2+^ imaging typically does not faithfully capture the Ca^2+^ concentration “seen” by the vesicular release site, even when using super-resolution techniques (e.g. ***Neef et al. (2018)***; ***Nakamura et al. (2015)***). Moreover, aside from this Ca^2+^ signal, transmitter release also depends the number of available SVs and / or release sites (***Neher, 2010***; ***Malagon et al., 2020***), and their docking or priming state (***Neher and Brose, 2018***; ***Taschenberger et al., 2016***). Finally, transmitter imaging offers insights into the spatiotemporal concentration profile of neurotransmitters, such as glutamate. One of the major tasks then is the interpretation of glutamate imaging data in the context of existing results acquired e.g. by electrophysiology and presynaptic Ca^2+^ imaging. For example, major efforts have been invested in the analysis of short-term synaptic plasticity by electrophysiology, yet, despite the great potential of neurotransmitter imaging for further elucidation of plasticity, much remains to be done to validate it against the electrophysiological benchmark.

### Virus-mediated expression of genetically encoded neurotransmitter indicators in the cochlear nucleus

Monitoring glutamate release with genetically encoded glutamate sensors requires a successful transfer of genetic material into the cells of interest. Here we chose to target SGNs and used cochlear injection of AAV to express iGluSnFR in ANF nerve terminals formed in the cochlear nucleus. For studying the cochlear nucleus, this approach has several advantages over other potential methods. First, compared to direct injections into the cochlear nucleus, strenuous craniotomy and potential damage of brain tissue can be avoided. Second, intracochlear injections target SGN predominantly, reducing transgene expression in primary cells in the cochlear nucleus. We reasoned that compared to virus-injection into the cochlear nucleus(e.g. ***Young and Neher (2009)***) expression limited to the presynaptic membrane of ANF would offer best specificity for analysis of afferent transmission to BCs. Previously, robust expression of iGluSnFR in SGN in the cochlea has been shown with an identical approach and was not associated with significant changes in auditory brainstem responses (***Özçete and Moser, 2021***).

There are multiple established options for introducing genetic material into the cochlea of living animals. First, embryonic otocysts can be manipulated either with the help of viral vectors (***Bedrosian et al., 2006***; ***Hernandez et al., 2014***) or electroporation (***Brigande et al., 2009***). Second, viral vectors can be injected into the cochlea of postnatal or adolescent mice (e.g. ***Akil et al. (2012)***; ***Jung et al. (2015)***; ***Keppeler et al. (2018)***; ***Akil et al. (2019)***). Postnatal cochlear AAV-mediated delivery proved very reliable: In injected mice, iGluSnFR expression in SGN nerve terminals in the cochlear nucleus could be confirmed by fluorescence microscopy and for almost all of them, successful measurements could be made.

Based on the confocal imaging of the stained injected cochleae, we assume that not all SGNs expressed iGluSnFR. Previous statistical analysis across several studies of AAV-mediated transgene expression of SGNs using various capsids but the same promoter (human synapsin) on average report a success rate of 70% (***Dieter et al., 2020***), although in a previous study transduction efficiency as high as 99% was described in the apical cochlea of at least one animal (***Özçete and Moser, 2021***). Incomplete transduction would explain our observation of a lack of iGluSnFR response to stimulation around some of the electrophysiologically and morphologically identified BCs, despite the presence of stereotypical eEPSCs. In the cochlea, we did not detect iGluSnFR-expression in the organ of Corti beyond the SGNs. In the cochlear nucleus, iGluSnFR immunofluorescence concentrated around cell bodies and co-localized to presynaptic active zones, often morphologically consistent with expression in endbulbs of Held. Neither in the cochlear nucleus nor in the spiral ganglion, systematic differences in expression based on tonotopy were obvious.

### Influence on physiological glutamatergic transmission

One important property of an ideal sensor for monitoring synaptic activity is that it does not alter physiological synaptic transmission. For GEGIs, there are multiple ways in which synaptic transmission may be affected. First, the surgery or the expression in neurons and glia may affect the viability and development of cells in which they are expressed by altering cellular processes. Second, even though iGluSnFR only has one C-terminal transmembrane domain, anchoring it in the plasma membrane (***Marvin et al., 2013***), the additional presence of protein in the presynaptic plasma membrane may affect the energetic balance of exocytosis and / or endocytosis by interaction with involved proteins or altering membrane curvature. A third possible way was suggested recently by ***Armbruster et al. (2020)***, highlighting the possibility of altered glutamate dynamics in the synaptic cleft, if additional glutamate-binding molecules (i.e. GEGI) are introduced. Such an effect is not implausible: iGluSnFR has an affinity to glutamate of around 84 µM (***Marvin et al., 2018***), and the affinity of AMPAR (specifically GluR3 and GluR4 subunits predominant in the BCs, see ***Gardner et al. (2001)***) to glutamate is also in the µM range (***Traynelis et al., 2010***), such that iGluSnFR could compete with AMPAR for glutamate molecules in the synaptic cleft. ***Armbruster et al. (2020)***, modelling single release sites, indicate that the effect of glutamate indicators might be largest ≥110 nm from the release site. The endbulb of Held is a large, calyceal synapse that employs tens to hundreds of active zones (***Nicol and Walmsley, 2002***; ***Neises et al., 1982***; ***Ryugo et al., 1996***, ***1997***; ***Butola et al., 2017***) and it has been suggested that synapses of a similar structure delay glutamate clearance for surprisingly long (***Otis et al., 1996***), influencing excitatory postsynaptic currents. Thus, we reasoned that a strong effect of iGluSnFR on physiological glutamate signalling would be particularly pronounced at this synapse.

In the electrophysiological data on hand, limited evidence of altered neurotransmission could be found. On the one hand, mEPSC decay and duration were found to be longer in postsynaptic bushy cells of iGluSnFR-expressing endbulbs, while on the other hand evoked responses were not significantly different. In order to detect more subtle putative effects, we also compared measures of release probability and apparent pool size.

We found a statistically significant increase in PPRs with a 10 ms but not with a 100 ms ISI at 2 mM [Ca^2+^]_*e*_. This observation would agree with a buffering effect, in which part of the glutamate released in the first stimulus is bound to iGluSnFR. As the fraction of bound iGluSnFR is supposedly still high after 10 ms (circa 48% of the initial number of bound iGluSnFR molecules, assuming monoexponential unbinding kinetics and a time constant of 13.8 ms (***Helassa et al., 2018***) a subsequent pulse leads to a comparatively larger postsynaptic response, as iGluSnFR copies present in the synaptic cleft are saturated with glutamate and thus unable to compete with AMPAR. At 100 ms, considerably fewer iGluSnFR copies are bound to glutamate (ca. 0.07% of the initial number given the same assumptions), thus the influence of a putative buffering effect is expected to be lower, which is consistent with the data. At 4 mM [Ca^2+^]_*e*_, no statistically significant change in PPRs could be observed. This may be seen as contradictory evidence to the data recorded at 2 mM [Ca^2+^]_*e*_. Another possible explanation for the difference in the data recorded at 2 mM [Ca^2+^]_*e*_ and 4 mM [Ca^2+^]_*e*_ could be that the relatively higher amounts of released glutamate at 4 mM [Ca^2+^]_*e*_ saturate either iGluSnFR or glutamate receptors, reducing the effect of additional glutamate buffering even in the presence of kynurenate. In this case, it would be worthwhile to explore differences in the effect of preand postsynaptic expression of iGluSnFR to see if iGluSnFR in the presynaptic plasma membrane more efficiently captures glutamate before it can reach postsynaptic receptors.

Our preliminary train stimulation analysis of vesicle pool dynamics in the presence and absence of AAV-mediated iGluSnFR expression in SGNs has not revealed significant differences between the two conditions. Further experiments, potentially involving faster versions of iGluSnFR and employing trains of different stimulation rates for model based analysis of vesicle pool dynamics (***Neher and Taschenberger, 2021***) will help to assess the value and impact of iGluSnFR in the analysis of transmission at calyceal synapses.

Previously, it was suggested that the introduction of iGluSnFR alters glutamate transporter currents in astroglia and at least our mEPSC recordings suggests that iGluSnFR presence indeed has a noticeable, albeit small effect on neurotransmission. The discrepancy between evoked and miniature EPSCs might not be too surprising, as eEPSCs represent summations of quantal events where some part of the variance is explained by differences in timing between quantal events. Thus, even slightly asynchronous summation of quantal events likely impacts the decay phase of the eEPSCs in addition to the deactivation kinetics of AMPAR currents (***Isaacson and Walmsley, 1995***; ***Diamond and Jahr, 1995***).

As it is generally unlikely that external manipulation of a complex, biological system has no effect on any process within the system, the more important question is whether the introduction of iGluSnFR, either by buffering glutamate in the synaptic cleft or by the expression system used, influences neurotransmission to a degree, which invalidates the obtained results.

The data on hand suggests that this is not the case. Firstly, even if a larger sample size may uncover more subtle effects neurotransmission of evoked events, our measurements suggest a small effect size. Secondly, even as we did find changes in mEPSC, it is probable that the biological significance is limited. The development of the auditory system is heavily influenced by spontaneous activity (***Wang and Bergles, 2015***; ***Babola et al., 2018***; ***Yu and Goodrich, 2014***) and iGluSnFR was introduced on postnatal day 6, during a developmentally vulnerable phase (***Tierney et al., 1997***). Nevertheless, our results give little indication of significantly impaired development, which agrees with previously published results of unaltered auditory brainstem responses of mice, which were subject to the same expression strategy as used here (***Özçete and Moser, 2021***).

Additionally, we cannot exclude the possibility that the surgery to deliver the viral vector to the inner ear or the expression system, rather than iGluSnFR itself, causes the subtle changes in synaptic transmission. Previously, a serotype-dependent effect of AAVs has been described in hippocampal synapses (***Jackman et al., 2014***). It would be interesting for future research to focus on the mechanism of the putative effect either by comparing transgenically to virally expressed iGluSnFR or comparing virally expressed iGluSnFR to another virally expressed fluorescent protein.

The, if at all, modest effect iGluSnFR expression has on synaptic transmission also stands in contrast to the rather pronounced effect GECIs may have on synaptic transmission, as e.g. presynaptic expression of GCaMP6m in the Calyx of Held strongly affects evoked release and synaptic plasticity (***Singh et al., 2018***).

### iGluSnFR response in the cochlear nucleus

Generally, the iGluSnFR response amplitude was large (Δ*F*∕*F*_0_ = 0.30 ± 0.02 at 2 mM [Ca^2+^]_*e*_), but rather variable between cells (see fig. 6 **A**). Since the method of selecting a ROI was dependent on large differences in Δ*F* between synapse and surrounding background, and cells for which a plausible ROI could not be determined were not analyzed further, cells included in the final dataset were selected to have a large Δ*F* .

Curiously, iGluSnFR recordings performed at 4 mM [Ca^2+^]_*e*_ did not have a higher amplitude peak than recordings performed at 2 mM [Ca^2+^]_*e*_, but instead increased glutamate release was reflected in longer decay times. Similarly, it has been reported before that stronger stimulation by multiple APs leads to a plateau of iGluSnFR signal intensity and to a slower decay (***Marvin et al., 2013***). Interestingly, within e.g. train data recorded at the respective extracellular calcium concentrations, clear graduations of amplitude could be observed, while decay appeared to stay stable.

Generally, we were able to resolve individual release events well under lower frequency stimulations, while peaks merged under higher frequency stimulation. Future imaging studies of glutamate release at calyceal synapses should explore the potential of new iGluSnFR variants with lower affinity that provide more rapid signal decay. This will ideally go along with imaging at higher framerate and might require stronger intensities of the excitation light to boost the fluorescence signal.

One important question in the interpretation of the iGluSnFR data is, which measure of the response best corresponds to the amount of glutamate that is released presynaptically.

The amplitude in response to a single stimulation was much more variable than the amplitude of eEPSCs. This discrepancy might be either due to differences in the postsynaptic sensitivity or due to differences in the fluorescent response to released glutamate and the limited variability of the recorded mEPSC amplitude suggests that the latter option is more likely. A potential difference in fluorescent response of two endbulbs to the same amount of released glutamate could be explained through multiple, plausible effects. Firstly, the imaging plane might be chosen better or worse for different endbulbs. Secondly, synapses may differ in their geometry and their orientation and synapses oriented along the *Z* axis may appear brighter than those oriented in the *X* − *Y* plane. Thirdly, correcting for baseline fluorescence might be insufficient to account for varying expression levels of iGluSnFR between cells especially if there is a large autofluorescent component or contribution of neighboring fluorescent ANFs.

Normalizing the area under the fluorescence curve and the corresponding eEPSC responses to trains of 5 or 10 stimuli at 10 Hz to the first response in the train increases the correlation between these two measures of release considerably. This observation is consistent with the idea that iGluSnFR responses are scaled differently for different endbulbs, but correlate within measured endbulbs with released glutamate.

In order to assess the relationship between iGluSnFR signal and electrophysiology more closely, we applied deconvolution analysis to the iGluSnFR data. For the deconvolution analysis, we assumed linearity of the system and immediate rise and exponential decay of the iGluSnFR signal. Even though the peaks in the deconvolved traces are relatively broad, suggesting additional components in the signal, the functional measures derived from deconvolution analysis still compared favorably to the electrophysiological measures. Deconvolving the iGluSnFR signals yields an even closer correlation between iGluSnFR and eEPSC signal. As AMPAR currents are thought to scale approximately linearly to released glutamate (assuming efficient pharmacological prevention of desensitization and saturation by 1 mM kynurenate), this linear relationship between iGluSnFR signal and eEPSCs suggests that iGluSnFR imaging is well-suited to obtain reliable information about presynaptic glutamate release under near-physiological conditions.

In this study we did not resolve glutamate signals arising from release of single SVs (i.e. quantal events). Optimizing imaging technique may reduce noise level, while the development of improved GEGIs could improve the signal to a level, at which spontaneous release events can be identified reliably in the cochlear nucleus. In retinal slices, where quantal events have been reliably observed with two-photon imaging, temporal deconvolution was successfully employed to estimate release rates from iGluSnFR signal (***Srivastava et al., 2022***; ***James et al., 2019***). Subcellular targeting of iGluSnFR variants to the postsynaptic membrane may reduce measurement errors introduced by contributing extrasynaptic iGluSnFR signal and improve spatial resolution of glutamate imaging data(***Hao et al., 2023***; ***Aggarwal et al., 2022***).

A second approach routinely used in electrophysiology studies is the calculation of quantal size based on the binomial statistics of evoked SV release. The application of this model to measured data assumes additionally that the variance of eEPSC size is determined by the variance given by the binomial statistics of SV release. Thus, parameters of the fit only approximate true values if the noise level in the recordings is low in comparison to the variance introduced by the stochastic nature of SV release. In common electrophysiological preparations, root-mean-square noise levels are routinely in the 100 fA range, while the standard deviation of eEPSC amplitude can easily be in the range hundreds of pA, yielding satisfactory results in the binomial analysis. In our case, variation in iGluSnFR amplitude was dominated by measurement noise, rendering this approach impractical. In previous studies, this approach was successfully implemented for EOS recordings under low measurement noise conditions in cell culture and with confocal microscopy (***Sakamoto et al., 2018***) and iGluSnFR in organotypic hippocampal cultures (***Dürst et al., 2022***). A related method based on the failure rate has been used for quantal analysis with Ca^2+^ imaging at smaller synapses in hippocampal slices (***Oertner et al., 2002***). Similarly, an approach based on the covariance of subsequent responses in a train of stimuli established in the calyx of Held (***Scheuss et al., 2002***) and also based on the binomial statistics of SV release, did not give meaningful results when applied to the data on hand (not shown).

In lack of satisfactory estimates for quantal size, the range of application of iGluSnFR in the presented conditions is limited, but important functional characteristics of synapses could be observed and quantified. For instance, we report a direct measurement of synaptic depression, which agrees well with electrophysiological control recordings. Specifically, deconvolution analysis of iGluSnFR signal closely tracked the recorded eEPSCs. Faithful measurement of presynaptic plasticity without the need for electrophysiological methods may enable future experiments, where short-term adaptations of large networks of neurons can be observed simultaneously.

Additionally, adapting standard methods from the analysis of cumulative eEPSCs (while considering the caveats summarized e.g. in ***Neher (2015)***), pool size can be determined as a multiple of the first response, but not in terms of individual vesicles.

## Methods and Materials

### Animal handling

We expressed iGluSnFR in SGNs of C57Bl/6J mice via viral vector injection into the neonatal cochlea as described previously (***Özçete and Moser, 2021***). Electrophysiological and imaging data was recorded from injected mice and their uninjected littermates of either sex from postnatal day 16 to 25 (P16 to P25). All experiments were performed in accordance to German national animal guidelines and were approved by the animal welfare office of the state of Lower Saxony as well as the local animal welfare office at the University Medical Center Göttingen (permit number: 17- 2394). The the AAV carrying the plasmid coding for iGluSnFR under the human synapsin promoter (pAAV.hSyn.iGluSnFR.WPRE.SV40) was a generous gift from Laren Looger (Addgene viral prep # 98929-AAV9) or produced in our own laboratory. Vector injections were performed as described before (***Jung et al., 2015***). In brief, P5 to P7 wildtype mice were anesthetized with isoflurane and the body temperature was kept at 37°C with a temperature plate and a rectal temperature probe. Under local anesthesia with xylocaine, the right ear was accessed through a dorsal incision and the round window exposed. 1 – 1.5 µL of suspended viral particles (titer 10^13^ vg/ml) were injected through the round window into the scala tympani through a quartz capillary. After the surgery, the incision wound was sutured and buprenorphine (0.1 mg/kg body weight) was applied for pain relief. The recovery status of the mice was monitored daily. Before and after the surgery, animals were kept with their mother and littermates until weaning (ca. P21) in a 12 h light/dark cycle, with access to food and water ad libitum.

### Tissue preparation

Brainstem slices were prepared as described previously (***Butola et al., 2017***). Briefly, mice were decapitated and the brain was rapidly removed and placed into ice-cold cutting solution, containing in mM: 50 NaCl, 120 sucrose, 20 glucose, 0.2 CaCl_2_, 6 MgCl_2_, 0.7 sodium ascorbate, 2 sodium pyruvate, 3 myo-inositol, 3 sodium l-lactate, 26 NaHCO_3_, 1.25 NaH_2_PO_4_ ⋅ H_2_O, 2.5 KCl with pH adjusted to 7.4 and an osmolarity of around 320 mOsm/l. Meninges were removed with a forceps and hemispheres were separated using a blade. After removing extraneous tissue, the brain stem was glued to a block with cyanoacrylate glue (Loctite 401, Henkel) and 150 µm thick parasaggital slices containing the cochlear nucleus were cut with a Leica VT1200 vibratome (Wetzlar, Germany) for imaging and electrophysiology. After cutting, slices were allowed to equilibrate at 35° C for 30 min in artificial cerebrospinal fluid (aCSF) and kept at room temperature until used for electrophysio logical recordings. The aCSF used for incubation contained (in mM: 125 NaCl, 13 Glucose, 1.5 CaCl_2_, 1 MgCl_2_, 0.7 sodium ascorbate, 2 sodium pyruvate, 3 myo-inositol, 3 sodium l-lactate, 26 NaHCO_3_, 1.25 NaH_2_ PO_4_⋅H_2_O, 2.5 KCl with pH adjusted to 7.4 and an osmolarity of around 310 mOsm/l. Slices were successively transferred to a recording chamber and continuously superfused with saline solution containing (in mM) 125 NaCl, 13 Glucose, 2 or 4 CaCl_2_, 1 MgCl_2_, 0.7 sodium ascorbate, 2 sodium pyruvate, 3 myo-inositol, 3 sodium l-lactate, 26 NaHCO_3_, 1.25 NaH_2_PO_4_ ⋅ H_2_O, 2.5 KCl with an osmolarity of around 310 mOsm/l and a pH of 7.4. The chamber was mounted on a Axioskop 2 FS plus upright microscope (Zeiss, Oberkochen, Germany) and the sample was illuminated with Differential interference contrast (DIC) optics through a 40 × / 0.8 NA water-immersion objective (Achroplan, Zeiss, Oberkochen, Germany).

### Electrophysiology

Electrophysiological recordings were obtained from BCs in the anteroventral cochlear nucleus, which was localized in its characteristic position in respect to cerebellum, the cochlear Nerve and the dorsal cochlear nucleus. Whole-cell patch clamp recordings of cells in the cochlear nucleus were performed with 2 – 4 MΩ pipettes pulled from borosilicate glass (GB150F, 0.86x1.50x80mm; Science Products, Hofheim, Germany) with a P-87 micropipette puller (Sutter Instruments Co., Novato, CA, USA, which were filled with intracellular solution, containing (in mM) 120 potassium gluconate, 10 HEPES, 8 EGTA, 10 sodium phosphocreatine, 4 ATP–Mg, 0.3 GTP–Na, 10 NaCl, 4.5 MgCl_2_, 0.001 QX-314 and 1 Alexa-596 anatomical dye, with an osmolarity of around 300 mOsm/l and a pH of 7.3 (adjusted with KOH). ANFs project onto multiple primary cells types in the cochlear nucleus (***Brawer et al., 1974***). In the AVCN, the two most prominent cell types are BCs and stellate cells (***Brawer et al., 1974***; ***Brawer and Morest, 1975***). These cells are distinct in morphology and physiology (***Wu and Oertel, 1984***) and can be differentiated in a number of ways. As identified in Golgi stains, BCs have one axon and a singular dendrite, while stellate cells have three or more protrusions (***Wu and Oertel, 1984***; ***Brawer et al., 1974***). Physiologically, stellate cells have a linear current-voltage relationship around the resting membrane potential and react to sustained depolarizing current injections with regularly firing action potentials (***Wu and Oertel, 1984***). Voltage clamped, they react to stimulation of the afferent ANF with relatively broad and small eEPSCs (***Wu and Oertel, 1984***) and in response to spontaneous release, they show infrequent, small, and broad mEPSCs (***Lu et al., 2007***). In contrast, the input resistance of BCs markedly decreases for depolarizing current injections and they show large and fast eEPSCs (***Wu and Oertel, 1984***) and mEPSCs (***Lu et al., 2007***). Additionally, ***Chanda and Xu-Friedman (2010)*** showed that stellate cells and BCs differ in response to two consecutive ANF stimulation and the ratio of the eEPSCs in response to two pulses, 10 ms apart, the 10 ms paired-pulse ratio (PPR), can be used to divide BCs and stellate cells in two non-overlapping clusters, as eEPSCs in stellate cell facilitate while they depress in BC. To obtain a uniform cell population and because QX-314 interferes with normal action potential generation and thus prevents the easiest and most reliable identification method in current-clamp, BCs identified in three ways: (i) through the frequency, size and decay time of their mEPSCs, once stimulation had been established through (ii) the response to consecutive stimulations and the width and duration of the eEPSCs and after finishing experiments and carefully retracting the patch pipette through (iii) morphological identification with fluorescence imaging Alexa-568 using a 585 nm LED (p100, CoolLED, Andover, UK) and a standard mCherry filter cube (Semrock, Rochester, NY, USA).

Data was recorded with a HEKA EPC 10 amplifier (HEKA Elektronik, Lambrecht/ Pfalz, Germany). Recordings were digitized at 40 kHz and low-pass filtered at 7.3 kHz. A liquid junction potential of 12 mV was corrected on-line. For confirmed BCs, EPSCs were evoked using monopolar direct current injections (5 – 20 µA), generated with a linear stimulus isolator (WPI Stimulus Isolator A365, World Precision Instruments, Sarasota, FL, USA). This was achieved by placing an electrode in a saline-filled patch pipette in a distance of 15 – 30 µm from the cell and during continuous monitoring of the membrane current, increasing the injected current until a single eEPSC could be triggered. The input was confirmed to be indeed monosynaptic by reducing the injected current until any further reduction lead to a complete failure of eEPSC generation. If successfully confirmed, the input was stimulated with breaks in between trains of stimuli of 30 s. All recordings were performed at-70 mV unless stated differently. mEPSC recordings were performed in 2 mM [Ca^2+^]_*e*_. eEPSCs were recorded in either 2 or 4 mM [Ca^2+^]_*e*_ and following drugs were added to the bath solution exclusively for eEPSC recordings: 10 µM bicuculline methchloride and 2 µM strychnine hydrochloride in order to block inhibitory GABAergic and glycinergic currents, respectively, and 1 mM sodium kynurenate to prevent AMPAR saturation and desensitization.

### Glutamate imaging

iGluSnFR was excited using a 470 nm LED (p100, CoolLED, Andover, UK) and glutamate signals were recorded using a sCMOS digital camera (OrcaFlash 4.0, Hamamatsu, Hamamatsu City, Japan) mounted to the microscope using astandard GFP filter cube (Semrock, Rochester, NY, USA). Optimal LED intensity was chosen empirically as the minimal intensity at which a response to 25 stimuli at 100 Hz could be seen reliably, usually at 0.5 – 1 mW/mm^2^.

Acquisition was triggered with a 5 mV pulse controlled by the patch clamp amplifier and synchronized to the current injections. For each stimulation, 1200 ms of iGluSnFR signal were recorded after the first stimulation. The prior 1300 ms were universally collected as background image.

### Immunohistochemistry

For immunohistochemical analysis, different routes were chosen with respect to the tissue at hand. For cochleae, blocks containing the entire inner ear were obtained from the same animals used for in-vitro electrophysiology and glutamate imaging and immediately fixated using 4 % (v/v) formaldehyde (FA) (diluted from 37% stock with phosphate-buffered saline (PBS)) for 20 min and washed with PBS. Cochleae were cryosectioned after a 0.12 mM EDTA decalcification. For brainstem, slices were fixed after electrophysiological experiments with 3% para-formaldehyde (PFA) for 15 min and washed with PBS. Sections were incubated for 1 h in goat serum dilution buffer (16% normal goat serum, 450 mM NaCl, 0.6% Triton X-100, 20 mM phosphate buffer, pH 7.4), primary antibodies were applied overnight at 4 °C, after which secondary antibodies were applied for 1 h at room temperature. The following antibodies were used for the staining of the cochleae: chicken anti-GFP (Abcam, 1:500), rabbit anti-calretinin (Synaptic Systems, 1:2000) and guinea pig anti-parvalbumin (Synaptic Systems, 1:300) and secondary AlexaFluor-labeled antibodies (goat anti-chicken 488 IgG, Thermo Fisher Scientific, 1:200; goat anti-guinea pig 568 IgG, Thermo Fisher Scientific, 1:200; goat anti-rabbit 633 IgG, Thermo Fisher Scientific, 1:200) For brainstem sections, primary antibodies used were an Alexa-488 labeled rabbit anti-GFP (Thermo Fisher Scientific, 1:400) and guinea pig anti-bassoon (Synaptic Systems, 1:500) and secondary goat anti-guinea pig 633 IgG (Thermo Fisher Scientific, 1:200). Confocal sections were acquired with a laser-scanning confocal microscope (Leica LSM780, Leica, Wetzlar, Germany), equipped with a 488 nm, a 561 nm and a 631 nm laser through a 63 × / 1.4 NA oil-immersion objective (Plan-Apochromat, Zeiss, Oberkochen, Germany). Pin hole size was set to 1.0 airy units.

### Data analysis

All electrophysiological data was exported from .dat files using Igor Pro 6.3.2 software (Wavemetrics, Lake Oswego, OR) running an instance of Patcher’ Power Tools (Abteilung für Membranbiophysik, Max-Planck-Institut für biophysikalische Chemie, Göttingen, Germany) to .ibw files.

For the analysis of mEPSCs, Igor Pro 6.3.2 software together with a customized instance of the Quanta Analysis procedure (***Mosharov, 2008***) were used. Data was smoothened with a binomial filter at a corner frequency of 8 kHz. Recording quality and low-frequency noise level were sufficiently good to preclude further notch filtering. Peaks were automatically detected in an additionally filtered trace (1-dimensional Gaussian filter with a corner frequency of 2 kHz) and an additionally filtered derivative trace (1-dimensional Gaussian filter, corner frequency of 4 kHz), using a 5σ cutoff and manually inspected to ensure correct classifications as event. The rising phase was fit with a simple affine function, while the decay phase was fit with a single exponential. For further analysis and illustration, the median fit parameters as well as an averaged wave was extracted per cell.

Other electrophysiological data was analyzed using a custom-written python script using the Neo wrapper for .ibw files (***Garcia et al., 2014***).

### Deconvolution analysis

For some parts of the analysis, timeseries data derived from glutamate imaging was subject to deconvolution analysis. Since different approaches have been used in imaging and electrophysiological studies, we applied two different methods to deconvolve iGluSnFR signal.

The first approach, a Wiener deconvolution algorithm based on the discrete Fourier transform and its inversion, has been used in electrophysiological studies in end-plate currents at the neuromuscular junction (***Van der Kloot, 1988***) and for image analysis of iGluSnFR data (***Taschenberger et al., 2016***; ***James et al., 2019***). The iGluSnFR signal *s*(*t*) was modeled as

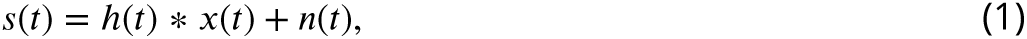

where ℎ(*t*) is the impulse response of the linear time-invariant system, *x*(*t*) the unknown signal, *n*(*t*) the additional noise and 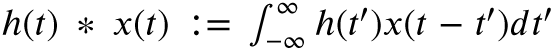 is the convolution of ℎ(*t*) and *x*(*t*). The Wiener deconvolution gives an estimate of *X*(*f*), the Fourier transform of *x*(*t*), as

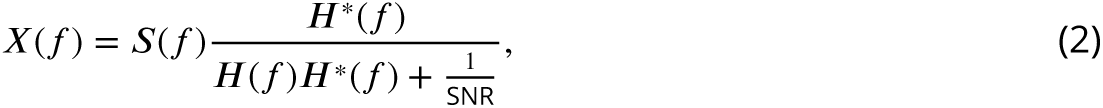

where *S*(*f*) is the Fourier transform of *s*(*t*), ***H***(*f*) is the Fourier transform of ℎ(*t*), ***H***^∗^(*f*) the complex conjugate of ***H***(*f*) and SNR the signal-to-noise ratio. The SNR was estimated as the squared quotient of maximum response amplitude and root-mean-square noise of the baseline of the recording. The deconvolution algorithm was implemented in a custom-written python script using the fast Fourier transform algorithm and matrix algebra tools provided in the numpy package.

A second method has been described for the analysis of end-plate currents at the neuromuscular junction (***Cohen et al., 1981***) and has recently been adapted to derive pool parameters from timeseries data generated by EOS imaging (***Sakamoto et al., 2018***). Here, a derivation for glutamate imaging data is presented.

Suppose that the glutamate content in one SV is able to induce a response of *n* iGluSnFR molecules, leading a change in fluorescence of *s*_0_ ∶= *n*⋅*g*_0_, where *g*_0_ is the unitary response. Between *t*^′^_1_ and *t*^′^_0_, in an interval *δt*^′^ = *t*^′^_1_ −*t*^′^_0_, SVs are released as a function of time *x*(*t*^′^_0_) *δt*^′^, each leading to a change in fluorescence of *s*_0_. Suppose further that the fluorescence is constant for each iGluSnFR copy over the course of *δt*^′^ and that iGluSnFR switch to a state of lower fluorescence stochastically and the time each iGluSnFR stays in the activated state is distributed exponentially, assuming monoexponential unbinding kinetics (***Helassa et al., 2018***).

The average iGluSnFR response *δs* during *δt*^′^ is thus given by

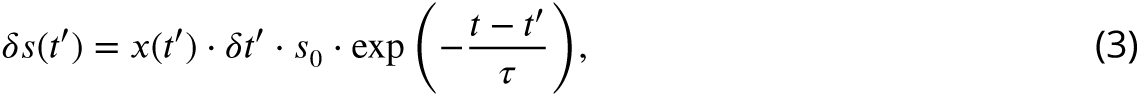

where τ is the average time an iGluSnFR copy stays in an activated state. Integrating equation 3 over all intervals yields

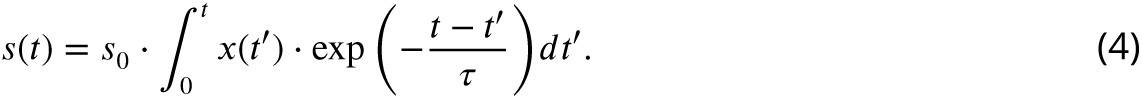

Differentiation with respect to *t* as a parameter and integration limit yields

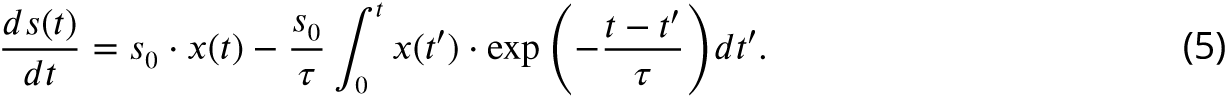

Substituting equation 4 reduces the expression to

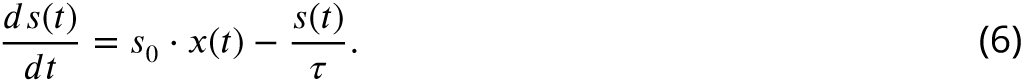

Thus, the signal can be deconvolved by

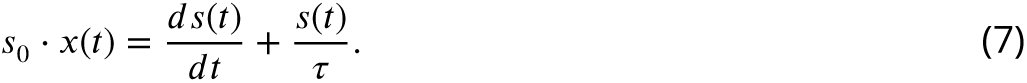

For computation, the derivative *ds*(*t*)∕*dt*(*t*^′^) was approximated as *s*(*t*^′^ + Δ*t*) − *s*(*t*^′^)∕Δ*t*. The error associated with *r* ∶= *s*_0_ ⋅ *x*(*t*) was approximated by

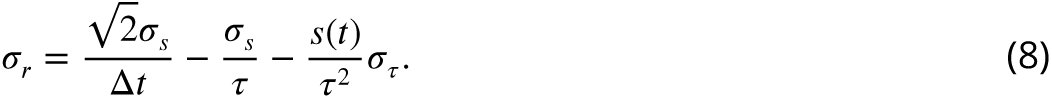

In electrophysiological recordings, quantal currents (either in the sense of unitary ion channel currents or currents induced by one released SV) are usually accessible through noise analysis. For glutamate imaging data, this approach has been successful in low-noise recordings of hippocampal cultures (***Sakamoto et al., 2018***). If the amplitude of the signal induced by one SV is not known, this approach still gives relevant information of relative release rates, e.g. over the course of multiple stimulations.

During stimulations, τ can be estimated as the time constant of a single exponential fit through the decay phase of the signal. This estimation is an upper bound for τ, as it assumes that all SVs are released and all iGluSnFR copies are activated simultaneously.

For practical calculations, this method was implemented in python, using a numerical approxi-mation to the derivative provided in the numpy package.

### Statistical analysis

If not indicated differently, means are given ± standard error of the mean. For derived quantities, standard errors were calculated taking gaussian error propagation into account. Statistical analysis was performed with consideration of clustering effects introduced by repeated measurements of the same cell, when appropriate. If clustering was not relevant to the statistical analysis, statistical significance was determined by the Wilcoxon rank sum test. When clustering had to be taken into account, condition effects were estimated using a mixed effects model with a random cell-specific intercept (for discussions of different approaches to clustered data in neuroscience research see ***Galbraith et al. (2010)***; ***Yu et al. (2022)***). When given, *p* values are presented as an exact number, except if smaller than 0.001 (for a discussion see ***Wasserstein and Lazar (2016)***). If effects are described as “statistically significant", a canonical significance level of *α* = 0.05 was assumed. When a number *m* of multiple comparisons were performed, Bonferroni-adjusted significance levels *α*^∗^ = *α*∕*m* are noted.

Statistical analysis was performed using custom-written python scripts, using implementations in the statsmodels and scipy packages.

## Acknowledgments

We thank Dr. Ö.D. Özçete for introducing PJHH to glutamate imaging and sharing preliminary image analysis scripts with us, Dr. E. Neher for helpful comments on data analysis and experimental design and N.I. Bosse for helpful comments on statistical analysis. We thank Ö.D. Özçete and Theocharis Alvanos for comments on the manuscript. We thank S. Gerke, D. Gerke, and C. Senger Freitag for expert technical assistance. This work was supported by the Deutsche Forschungsge meinschaft (DFG, German Research Foundation) under Germany’s Excellence Strategy—EXC 2067/1-390729940 to T.M. as well as via the collaborative research center 1286 to T.M..

---

**Figure 1—figure supplement 1.**
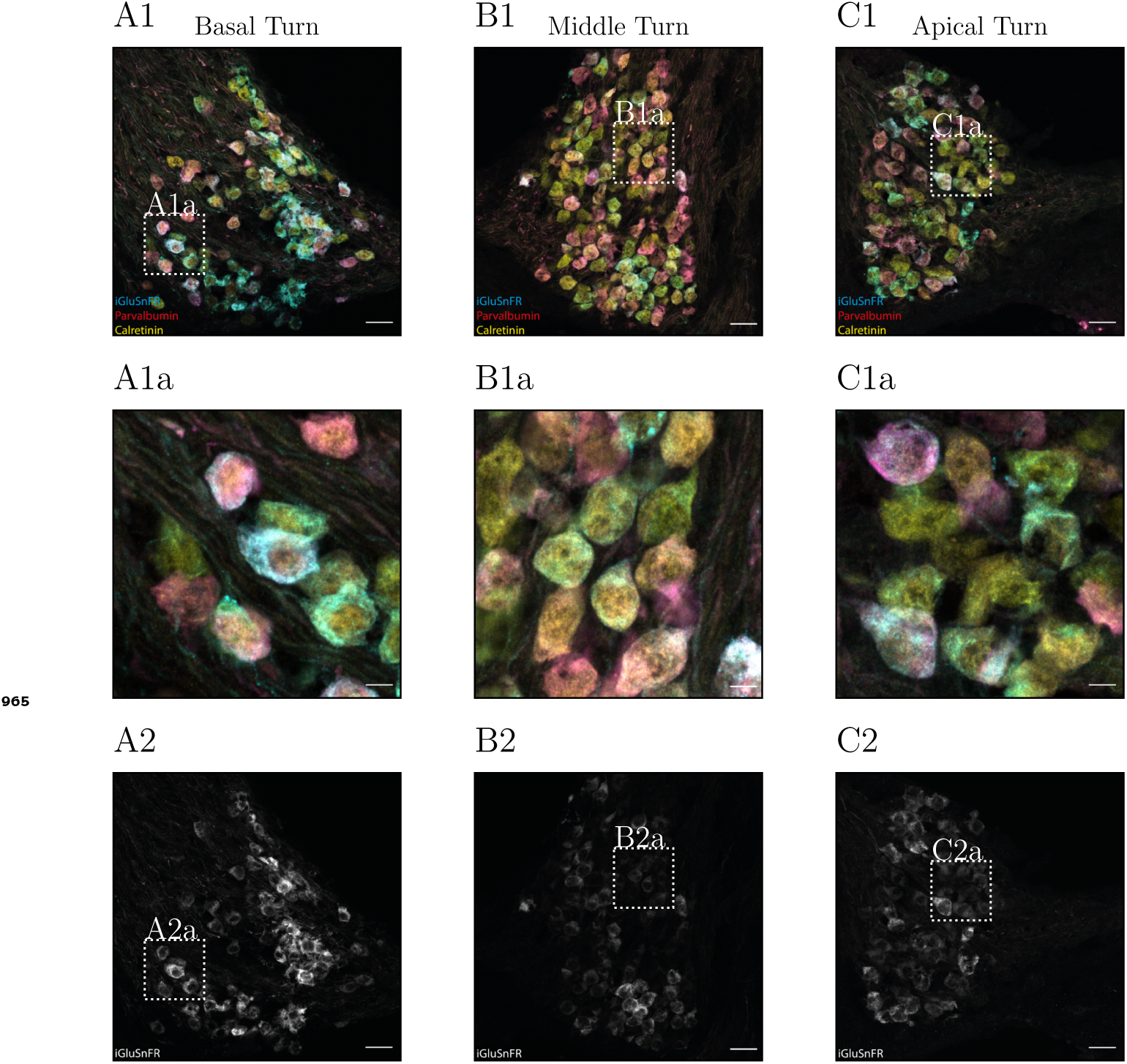
Immunohistochemistry of the cochlea. Confocal sections of cryosections of the spiral ganglion of a mouse cochlea, 11 days after viral transfection. GFP / iGluSnFR were stained using a primary antibody conjugated with Alexa-488, Parvalbumin with a secondary antibody conjugated with Alexa-561 and Calretinin with a secondary antibody conjugated with Alexa-633. In A, sections through the basal turn, in B trough the middle turn and in C through the apical turn of the cochlea are shown. In 1, iGluSnFR (cyan), Parvalbumin (red) and Calretinin (yellow) are stained. In 1a, details in 4 × zoom are shown. The respective regions are indicated by white squares in 1 (cont.)

**Figure 1—figure supplement 2.**
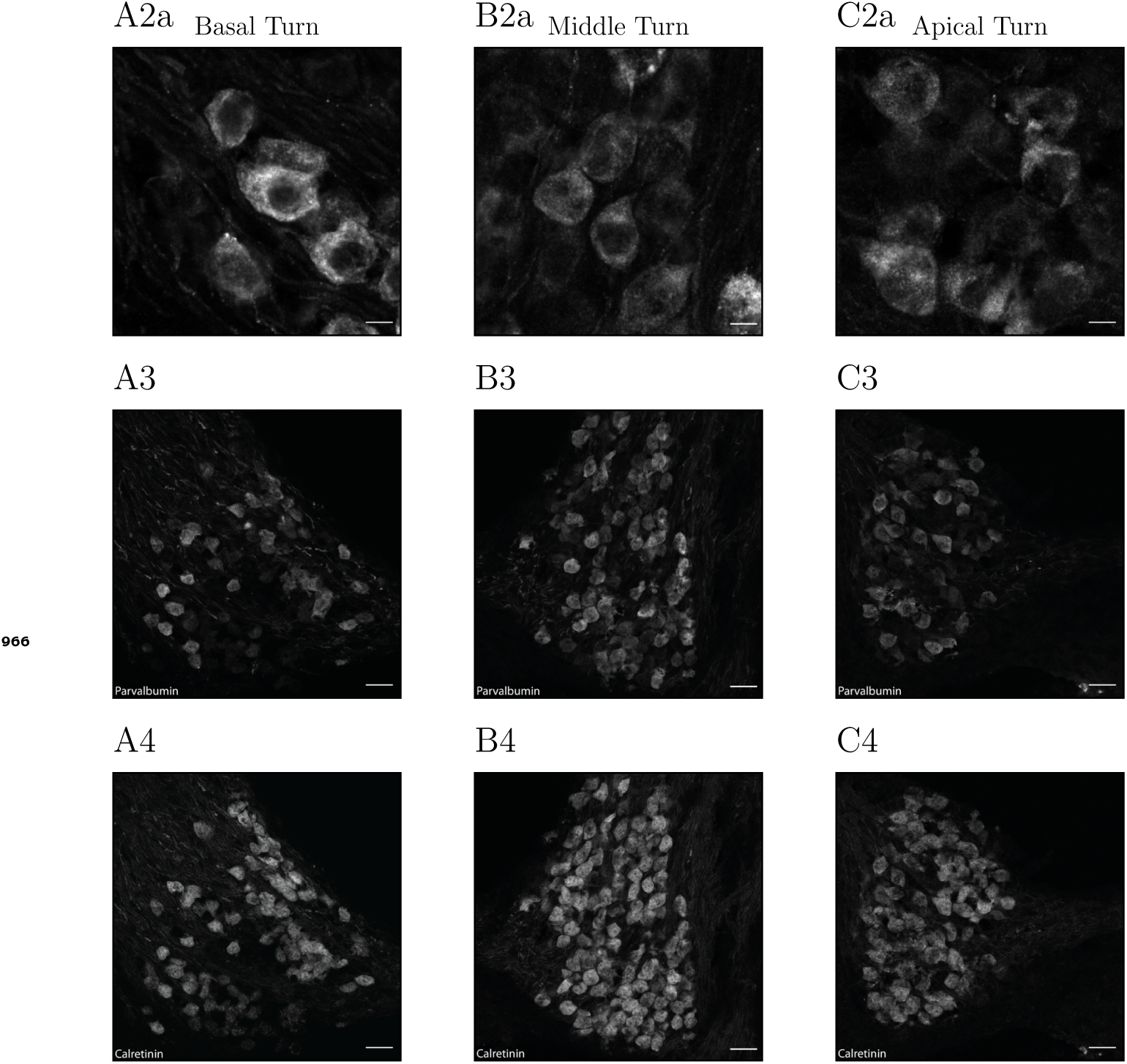
Immunohistochemistry of the cochlea (cont.) Grayscale images of the 488 nm / iGluSnFR channel (2), the 561 nm / Parvalbumin channel (3) and the 633 nm / Calretinin channel (4) are shown in the respective panels. For the 488 nm / iGluSnFR channel, in 4 × magnifications, 2a details are shown in the respective panels. Scale bars in the bottom right corner indicate 20 µm in all sections, except for the magnifications, where they represent 5 µm.

**Figure 1—figure supplement 3.**
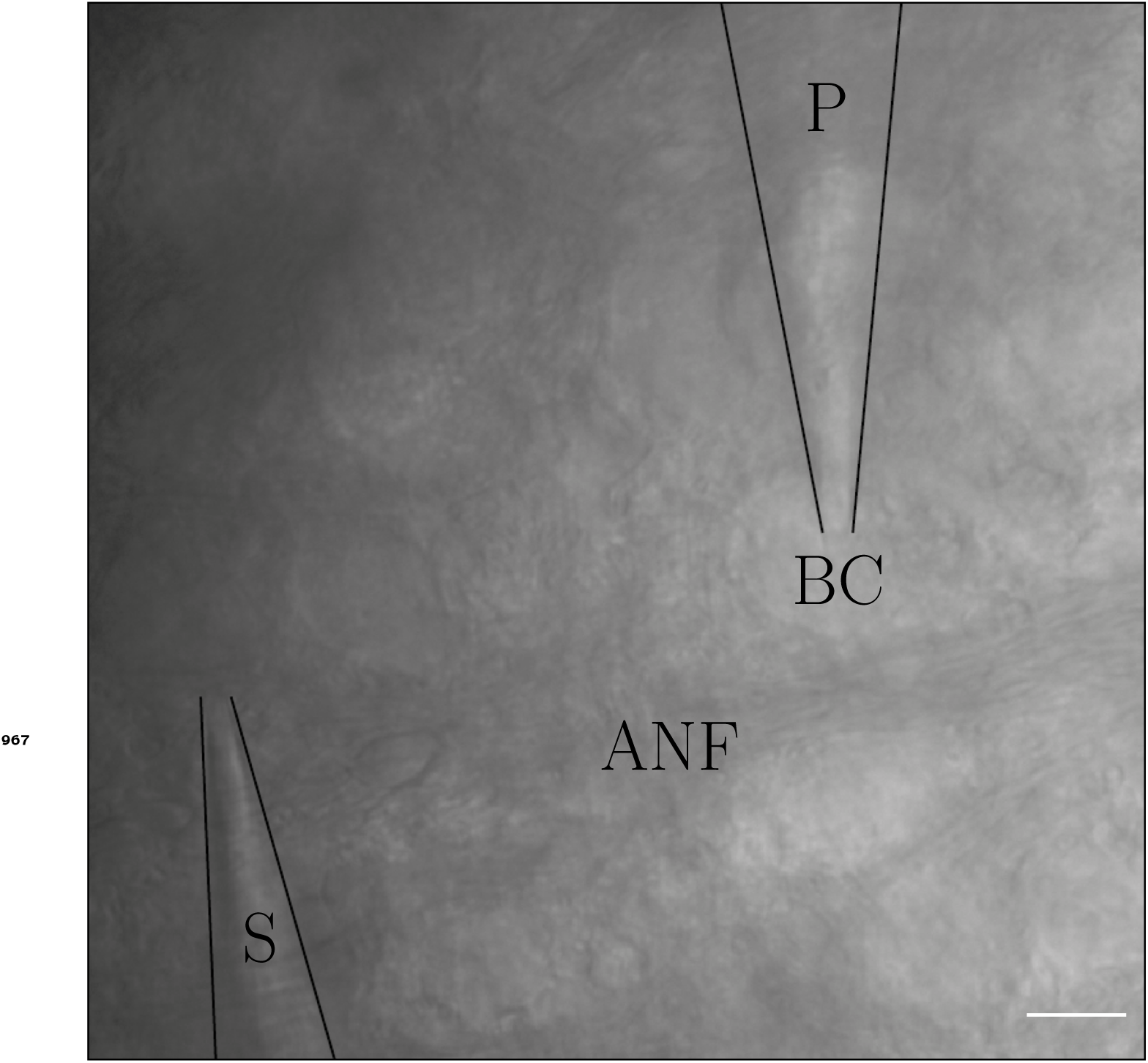
Bright-field example of a typical recording situation. DIC image of the experimental setup in a 150 µm slice of the cochlear nucleus. Here, the conditions during recording are shown. A postsynaptic bushy cell (BC) is accessed and voltage-clamped through a patch pipette (P), while simultaneously stimulated via a stimulation electrode, located in a saline-filled pipette (tip marked as S). In this image, the auditory nerve fibers (ANF) which transmit the stimulation signal to the presynaptic axosomatic terminal can be identified. Successful stimulation was, however, not dependent on successful identification of a fiber bundle leading directly to the postsynaptic BC, but was usually achieved by moving the stimulation electrode around the BC, while periodically injecting current in the tissue and monitoring the postsynaptic currents at the same time. Scale bar represents 20 µm.

**Figure 1—figure supplement 4.**
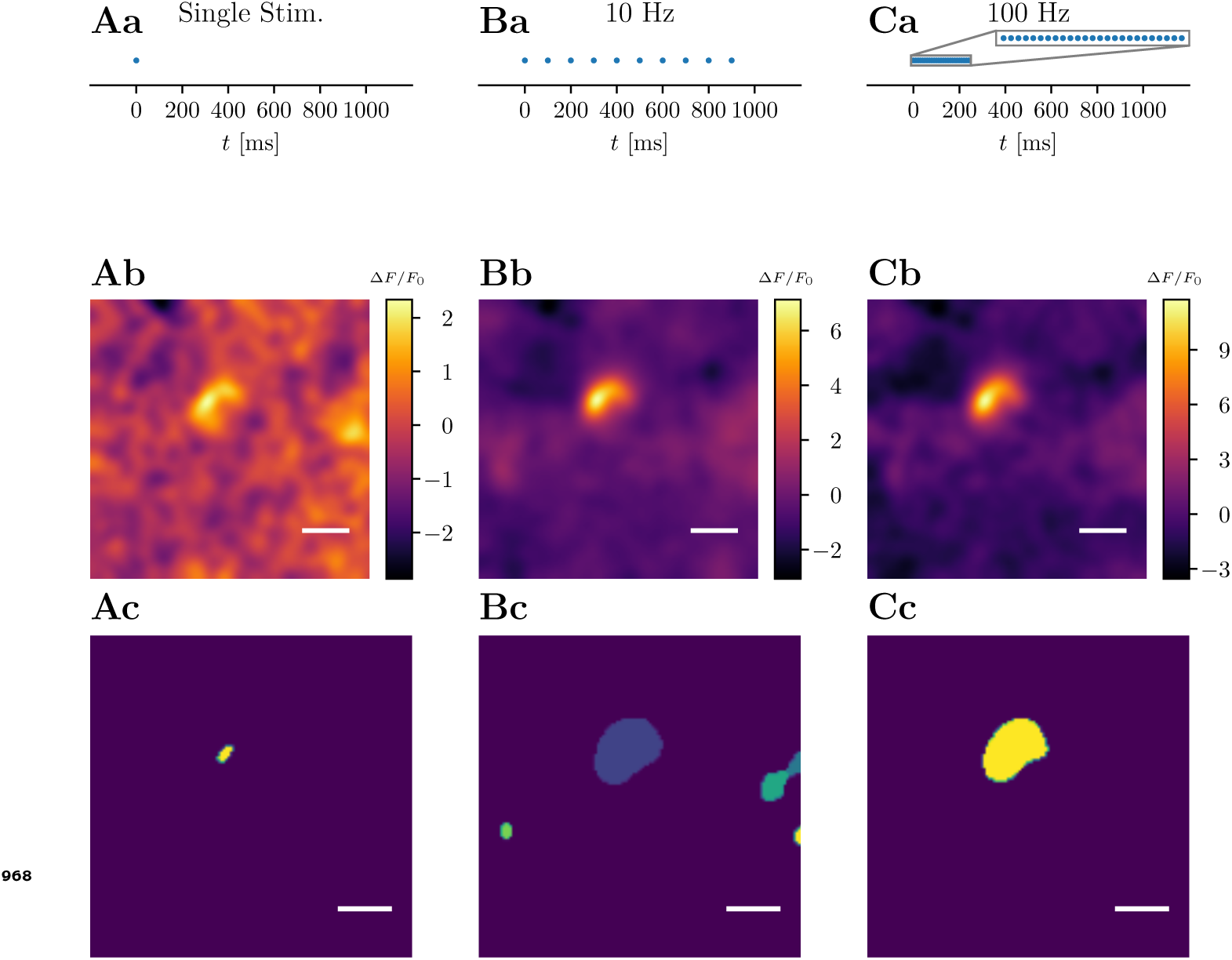
Comparison of ROI identification at different stimulation paradigms. While single stimulation leads to relatively noisy Δ*F* images, stimulation with 10 stimuli at 10 Hz or 25 stimuli at 100 Hz led to a Δ*F* image, which resembled terminal morphology more closely and allowed extraction of a meaningful ROI with minimal manual reevaluation. For this figure and further analysis, multiple recordings (usually 3 – 10) for each condition were averaged on a pixel-by-pixel basis. A: Single stimulation. Panel a: Scheme of stimulation paradigm. Filled dots represent a single afferent fiber stimulation. Panel b: Δ*F* image showing the relative average change in fluorescence of a single pixel from the beginning of the stimulation until ∼ 50 ms after the stimulation. Panel c: Automatic ROI detection without manual adjustments. B: Low-frequency stimulation of 10 stimuli at 10 Hz. Panel a: Scheme of stimulation paradigm in panel b: Δ*F* image showing the relative average change from the beginning of the stimulation until ∼1000 ms after the stimulation. Panel c: Automatic ROI detection without manual adjustments. C: High-frequency stimulation with 25 stimuli at 100 Hz. Panel a: Scheme of stimulation paradigm. Panel b: Δ*F* image showing the relative average change from the beginning of the stimulation until ∼ 390 ms after the stimulation. Panel c: Automatic ROI detection without manual adjustments. Small detected segments occurring outside of the central, labeled segment were removed manually afterwards, if they occurred, as they most likely either represent imaging noise or terminals projecting onto neighboring cells. Scale bars represent 5 µm.

**Figure 4—figure supplement 1.**
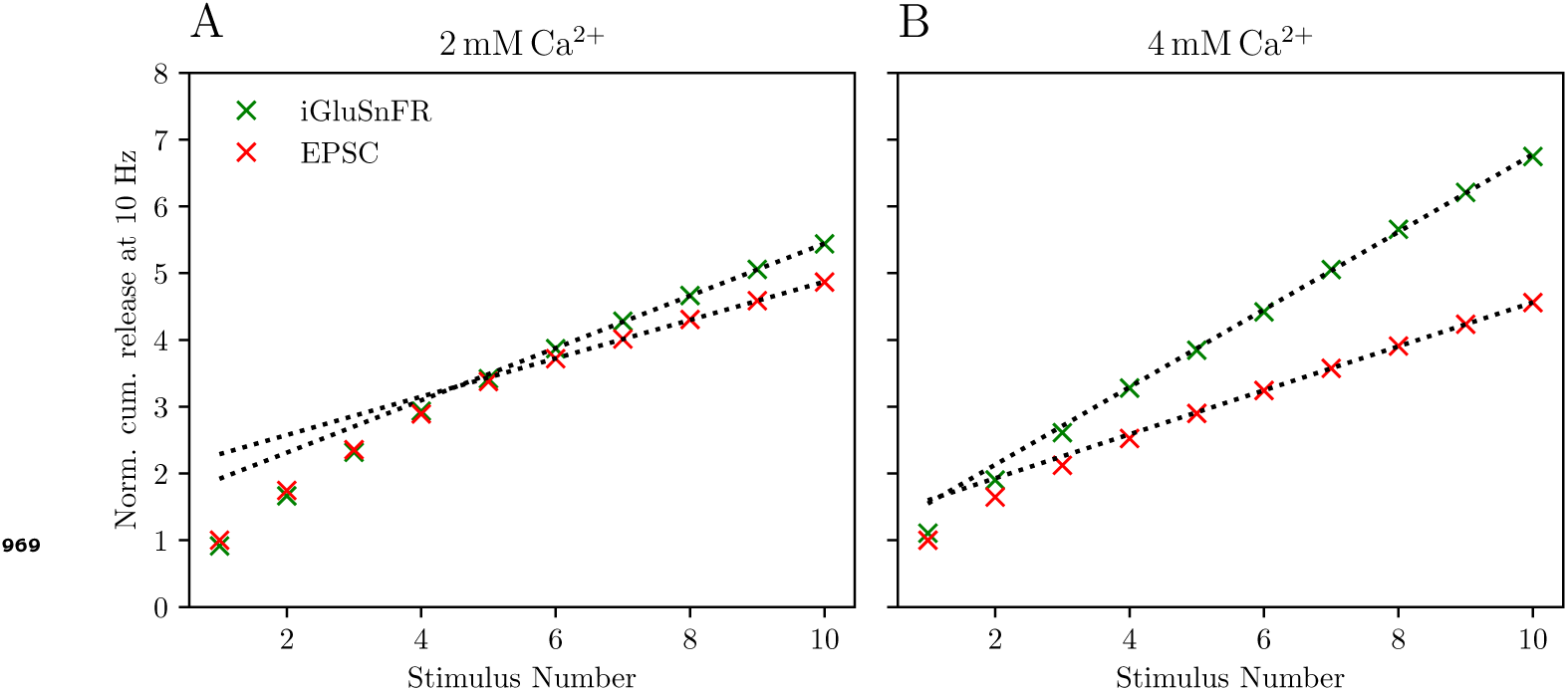
Cumulative release analysis of 10 Hz data. Cumulative release was derived by summation of peak amplitudes of consecutive events in electrophysiological data and deconvolved iGluSnFR signal (in this case of the 10 Hz data). Cumulative release was normalized by dividing by the single response at the respective [Ca^2+^]_*e*_ levels. Panel A show the normalized average cumulative release, derived from measurements of *N* = 7 animals, *n* = 8 cells, when stimulated with 5 or 10 stimuli at 10 Hz at 2 mM [Ca^2+^]_*e*_. Panels B show the average normalized cumulative release from *N* = 7 animals, *n* = 9 cells, when stimulated with 10 stimuli at 10 Hz at 4 mM [Ca^2+^]_*e*_.

**Figure 6—figure supplement 1.**
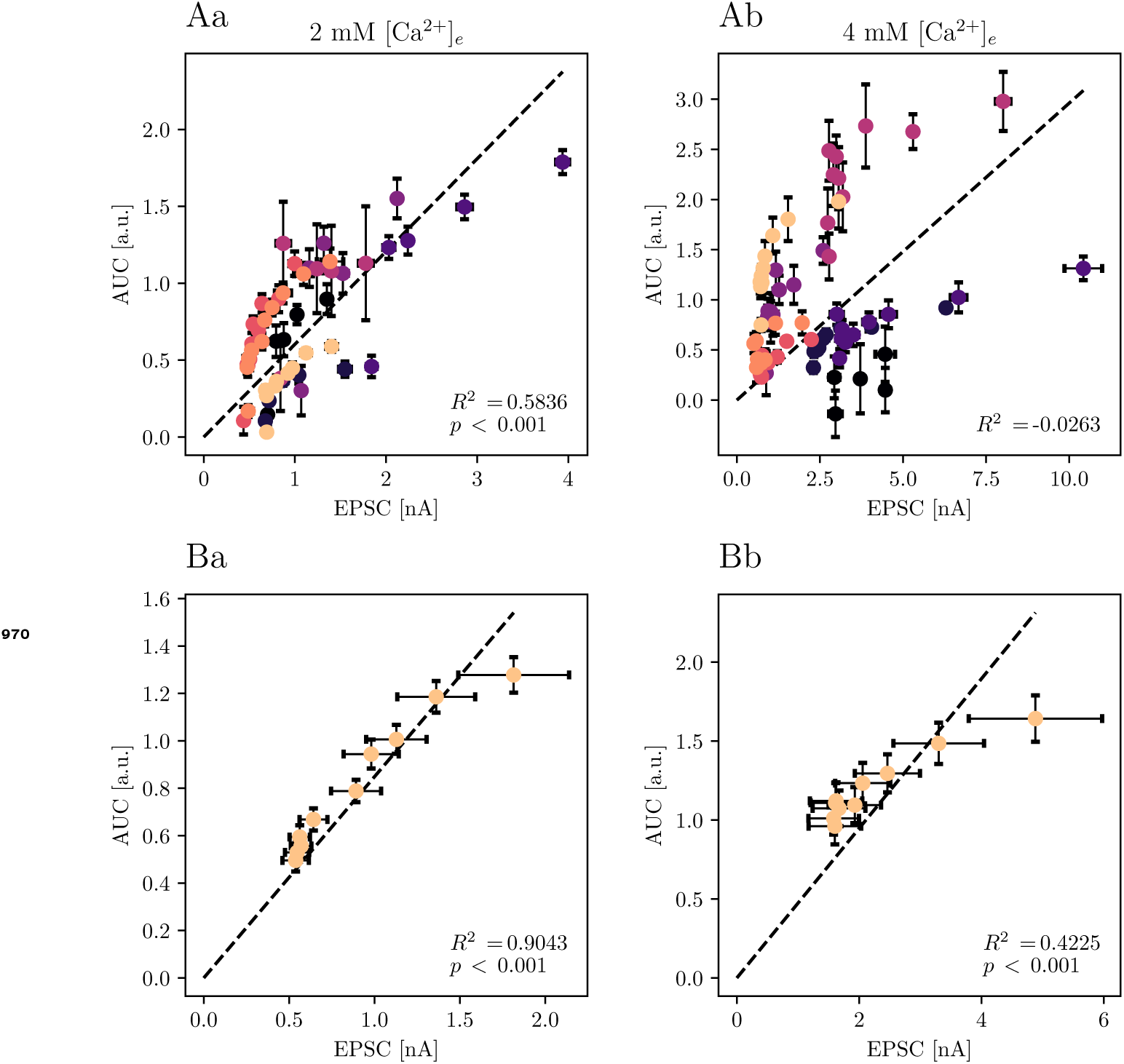
Non-normalized data of iGluSnFR vs. eEPSC data. Plots in A show the absolute amplitudes of AUC and eEPSC of the same data presented in fig. 6, **A**. Plots in B show the absolute amplitudes of AUC and eEPSC of the same data presented in fig. 6, **B**. The non-normalized data for 4 mM [Ca^2+^]_*e*_ was not fit well by a single linear function, likely due to larger variance between cells.

**Figure 6—figure supplement 2.**
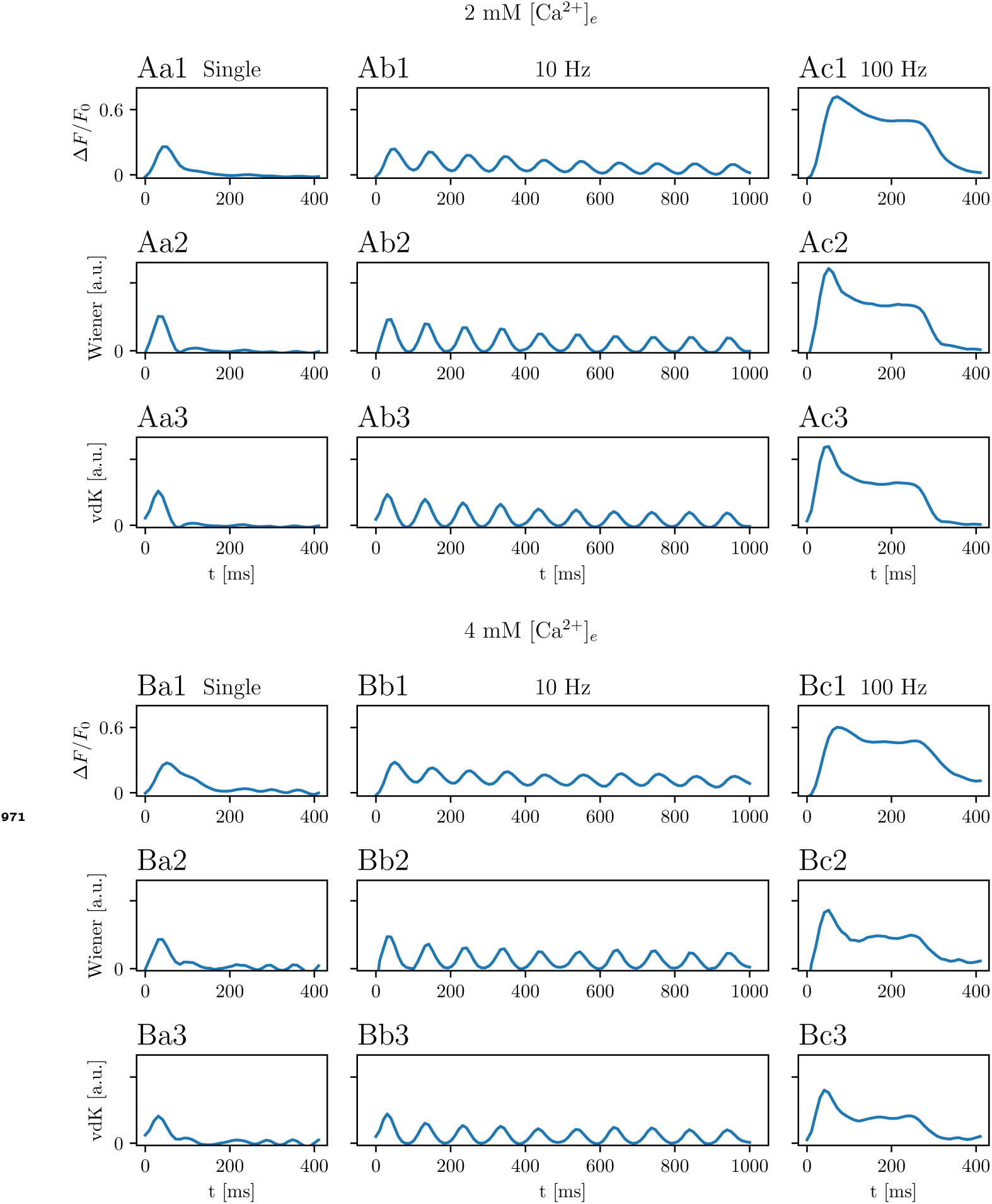
Deconvolved traces of averaged recordings. Recordings of *N* = 7 animals, *n* = 8 cells at 2 mM [Ca^2+^]_*e*_ (panels A) and *N* = 7 animals, *n* = 9 cells at 4 mM [Ca^2+^]_*e*_ (panels B) were averaged and filtered (panels 1). Either a Wiener deconvolution algorithm (panels 2, "Wiener") or a deconvolution algorithm based on ***Cohen et al. (1981)*** (panels 3, "vdK") was applied to the traces with single stimuli (panels a), 10 Hz traces (panels b) or 100 Hz traces (panels c). Both methods have been used (in variations) in the analysis of electrophysiological (***Cohen et al., 1981***; ***Van der Kloot, 1988***), as well as glutamate imaging data (***Taschenberger et al., 2016***; ***Sakamoto et al., 2018***). They both share the assumptions that (a) the iGluSnFR signal rises instantaneously, (b) glutamate release occurs completely synchronized and (c) iGluSnFR responses are added linearly and are indifferent to the history of responses. At our hands, results with both methods were roughly comparable. For further analysis, the analytical solution was used, as it has been proven to be more reliable for deconvolving electrophysiological responses (***Van der Kloot, 1988***).

## Notes

### Competing Interest Statement

The authors have declared no competing interest.

### Summary of Updates

- We initially reported data as mean +- s.e.m. over all data points, which has led to misunderstandings. We now report mean +- s.e.m. over the cell median. Additionally, we added captions to the tables to describe the statistical methodology in more detail. - We are unable to resolve the size of the readily releasable pool of synaptic vesicles in terms of individual vesicles. We modified the wording of our text to reflect this limitation. We updated the data presentation in figure 3 to be more easily understandable. We increased the text size in most figures. - Our attention was directed to a recent study probing synaptic transmission at the retina. We now discuss this study. - Our attention was directed to recently developed postsynaptically targeted iGluSnFR variants. We now discuss the relevant studies. - We fixed some typos, unclear wordings, superfluous abbreviations, unit conversion errors, and referencing issues.

